# Optical coherence tomography enables longitudinal evaluation of cell graft-directed remodeling in stroke lesions

**DOI:** 10.1101/2024.10.09.617387

**Authors:** Honour O Adewumi, Matthew G Simkulet, Gülce Küreli, John T Giblin, Arnaldo Bisbal Lopez, Şefik Evren Erdener, John Jiang, David A Boas, Timothy M O’Shea

**Affiliations:** Department of Biomedical Engineering, Boston University, Boston, MA, 02215-2407, USA; Institute of Neurological Sciences and Psychiatry, Hacettepe University, Ankara, 06230, Türkiye

## Abstract

Stem cell grafting can promote glial repair of adult stroke injuries during the subacute wound healing phase, but graft survival and glial repair outcomes are perturbed by lesion severity and mode of injury. To better understand how stroke lesion environments alter the functions of cell grafts, we employed optical coherence tomography (OCT) to longitudinally image mouse cortical photothrombotic ischemic strokes treated with allogeneic neural progenitor cell (NPC) grafts. OCT angiography, signal intensity, and signal decay resulting from optical scattering were assessed at multiple timepoints across two weeks in mice receiving an NPC graft or an injection of saline at two days after stroke. OCT scattering information revealed pronounced axial lesion contraction that naturally occurred throughout the subacute wound healing phase that was not modified by either NPC or saline treatment. By analyzing OCT signal intensity along the coronal plane, we observed dramatic contraction of the cortex away from the imaging window in the first week after stroke which impaired conventional OCT angiography but which enabled the detection of NPC graft-induced glial repair. There was moderate, but variable, NPC graft survival at photothrombotic strokes at two weeks which was inversely correlated with acute stroke lesion sizes as measured by OCT prior to treatment, suggesting a prognostic role for OCT imaging and reinforcing the dominant effect of lesion size and severity on graft outcome. Overall, our findings demonstrate the utility of OCT imaging for both tracking and predicting natural and treatment-directed changes in ischemic stroke lesion cores.

## Introduction

Stroke injuries are among the leading causes of death and disability worldwide and indiscriminately affect people across all age, ethnic, and racial groups (Feigin et al., 2017; Prust et al., 2024). Ischemic strokes caused by brain vessel occlusions, account for most clinical strokes (Phipps and Cronin, 2020). More severe hemorrhagic strokes involving vessel rupture, make up 10-15% of cases (Grysiewicz et al., 2008); however, ischemic strokes can undergo a hemorrhagic transformation in as many as 40% of patients which can increase morbidity (Jickling et al., 2014). An aging population has seen rising numbers of individuals experiencing recurrent, asymptomatic white matter strokes (Sozmen et al., 2012). These strokes can progress to larger injuries or undergo hemorrhagic transformations resulting in cognitive decline, motor impairments, or death with limited effective interventions (Rosenzweig and Carmichael, 2013; Smith et al., 2017). Clinical therapies for stroke are limited to acute thrombolysis (e.g. alteplase) or endovascular thrombectomy, but can only be applied to ischemic injuries within a narrow temporal window after stroke onset (Hacke et al., 2008), and many stroke patients cannot receive such therapies for logistical and socioeconomic reasons (Steven R. Messé et al., 2016) or because of a high risk of hemorrhagic transformation (Goncalves et al., 2022; Yaghi et al., 2017). There are no effective interventions for hemorrhagic stroke beyond surgery and hemostatic agents that stem bleeding at damaged vessels (Magid-Bernstein et al., 2022). While rehabilitation paradigms can aid in stimulating the plasticity of surviving neural circuits to promote modest functional recovery after stroke (Johansson, 2000), there are currently no therapies that can regenerate the regions of neural tissue damaged by stroke (Zhao and Willing, 2018).

Infarct- or hemorrhage-induced injury causes irreversible damage to neural tissue resulting in the formation of a stroke lesion core. Natural functional recovery after stroke depends on a combination of local neural circuit remodeling and neurogenesis directed by migrating neuroblasts from the subventricular zone (SVZ) (Liang et al., 2019). Significant natural recovery only occurs in regions of preserved neural tissue adjacent to stroke lesion cores because it contains essential glial cells that serve important support functions (Liang et al., 2019). At stroke lesion cores, damaged neural tissue is replaced by non-neural fibrotic scar tissue which is permanently devoid of these essential glial cells, so there is no support for spontaneous remodeling or regeneration of neural circuits at stroke lesion cores (Dias et al., 2021; O’Shea et al., 2020). Therefore, in large ischemic and hemorrhagic stroke lesions, post-stroke neurogenesis and neural circuit remodeling is limited, and natural functional recovery is poor. Strategies to alter the formation, size, or composition of non-neural lesion cores could augment post-stroke neurogenesis and/or neural circuit remodeling leading to improved recovery outcomes.

In certain brain injuries in the mammalian neonate, scar-free repair coordinated by immature astrocytes and microglia provides a supportive cell framework for neurogenesis, axon regeneration, and neural circuit remodeling (Berry et al., 1983; Li et al., 2020; Maxwell et al., 1990). This glial repair competency is lost in adult injuries (Li et al., 2020). Using cell grafts to provide exogenous sources of cells capable of glial repair of adult stroke injuries during the acute injury period beyond the thrombolytic therapy window (1-5 days post-injury) may be a functional basis for rebuilding neural tissue at stroke lesion cores. Recently, we demonstrated that neural progenitor cells (NPC) grafted into mouse ischemic striatal strokes at 2 days after injury spontaneously differentiate into astrocytes that direct glial repair at stroke lesions and serve neuroprotective functions such as restricting inflammation and preventing fibrotic scarring (O’Shea et al., 2022). However, NPC grafted into larger ischemic and hemorrhagic stroke lesions are less effective at remodeling lesion cores because cell fate and glial repair functions are strongly influenced by non-cell autonomous cues from lesion environments (Adewumi et al., 2024). Because stroke lesion cores evolve temporally, understanding how lesion environments alter the functions of cell grafts after transplantation requires being able to monitor lesion phenotypes longitudinally. Additionally, since lesion states influence grafting outcomes so significantly, it is important to understand, prior to treatment, whether a particular lesion phenotype is suitable for a grafting intervention and identify the time post-injury that would be most appropriate for cell grafting in order to optimize outcomes from these therapies.

There have been substantial advances in our understanding of the multicellular interactions involved in CNS wound responses after CNS injury through the advent of transgenic mouse models and immunohistochemistry methods (Burda and Sofroniew, 2014). However, most insights in this area have come from post-hoc evaluations of fixed brain sections which only provide information at a single terminal time point. These methods do not afford a way to track how lesions evolve over time in individual animals. Intravital imaging methods provide a viable way to temporally track dynamic, naturally occurring wound repair processes at stroke lesion cores, as well as how these processes can be altered by therapeutic interventions such as cell grafting. Two-photon microscopy has been used to evaluate naturally induced changes following ischemic stroke and reperfusion (Li and Murphy, 2008), but this imaging method requires the cells of interest to express fluorescent reporters in order to be identified and tracked. Optical coherence tomography (OCT), a non-invasive, label-free imaging modality, has recently become a popular and impactful neuroscience tool for characterizing cerebral blood flow, capillary dynamics and neural activity (Ruikang K. Wang et al., 2017; Tang et al., 2021). We and others have explored the use of OCT in rodent stroke models to track the size of stroke lesions acutely (Choi et al., 2019; Srinivasan et al., 2013; Sunil et al., 2021). Despite an incomplete mechanistic understanding, after ischemic stroke there is a well-documented increase in optical scattering in stroke lesion cores immediately after injury in rodent models as measured by OCT. Increased light scattering has been identified as a reliable predictor of stroke lesion core size at hyperacute (<4 hours) timepoints (Choi et al., 2019; Srinivasan et al., 2013) and acute timepoints, up to 72 hours after stroke (Sunil et al., 2021). However, while OCT has been used to identify infarcted tissue regions acutely after stroke and to evaluate vascular remodeling in the peri-infarct region several weeks after stroke, this intravital imaging technique has not been used to longitudinally track sub-acute wound repair processes or outcomes of experimental treatments, such as cell-grafting approaches, at stroke lesion cores.

We hypothesized that OCT could be an effective, label-free way to non-invasively identify and monitor natural wound repair processes at stroke lesion cores in mice and to study how a cell grafting intervention alters these natural wound repair outcomes. To advance the adoption of intravital imaging methods in preclinical stroke studies, we used OCT to longitudinally track changes in photothrombotic stroke lesion cores with and without experimental cell grafting interventions over the course of 2 weeks. We developed a new image processing and analysis framework to extend the interpretable data derived from OCT angiograms and identify new, temporally evolving features of the stroke lesions such as cortical atrophy and tissue layer separation due to lesion contraction. We correlated key OCT lesion signatures with cell and molecular features characterized by post-hoc immunohistochemistry. Cell grafting interventions applied at two days after photothrombotic stroke induction modestly altered the overall size and cellular phenotype of stroke lesion cores that enabled identification of cell graft-induced changes by OCT. Our findings demonstrate the feasibility of using OCT as a label-free method to monitor stroke wound repair processes and represent a promising tool for determining the suitability of individual stroke lesions for therapeutic interventions such as cell grafting.

## Results

### 1. Processing framework for interpreting OCT imaging of stroke lesions

We adapted previously developed OCT acquisition methodologies to longitudinally track wound healing at photothrombotic stroke lesions in mice during the sub-acute injury phase. To enable OCT image acquisition, we first performed a craniotomy over the cerebral cortex of the left hemisphere and installed a 3 mm diameter optical window two weeks prior to stroke induction. After acquiring baseline OCT images, we induced a photothrombotic stroke to a selected pial arterial vessel through the optical window, and OCT imaging was repeated one day after stroke to assess acute lesion phenotype. At two days post-stroke, mice underwent a second surgery to infuse 4 µL total of saline or neural progenitor cells (NPC) in media using four individual injections made in locations evenly spaced around the stroke. OCT images were acquired at one and two weeks post-stroke to characterize sub-acute wound healing processes, and mice were perfused immediately after the last OCT imaging session for post-hoc immunohistochemistry (IHC) evaluation (**Figure 1A**). Volumetric OCT images are generated through accumulating a series of A-lines, profiles of reflectivity over a defined tissue depth in the z-axis at a single pixel in the x-y plane (**Figure 1B**). B-scans are a rastered row of several A-lines, and C-scans are a series of accumulated B-scans representing an image volume. Raw reflectivity (RR) is used here to represent the acquired data within a single B-scan taken at a single defined timepoint. To extract key stroke lesion parameters, we optimized a series of different processing steps on the acquired OCT data to generate: (i) OCT angiograms, (ii) Coronal profiles, and (iii) OCT scattering maps. The processing required to derive these different forms of OCT information from acquired OCT scans warrants brief overview.

**Figure 1:**
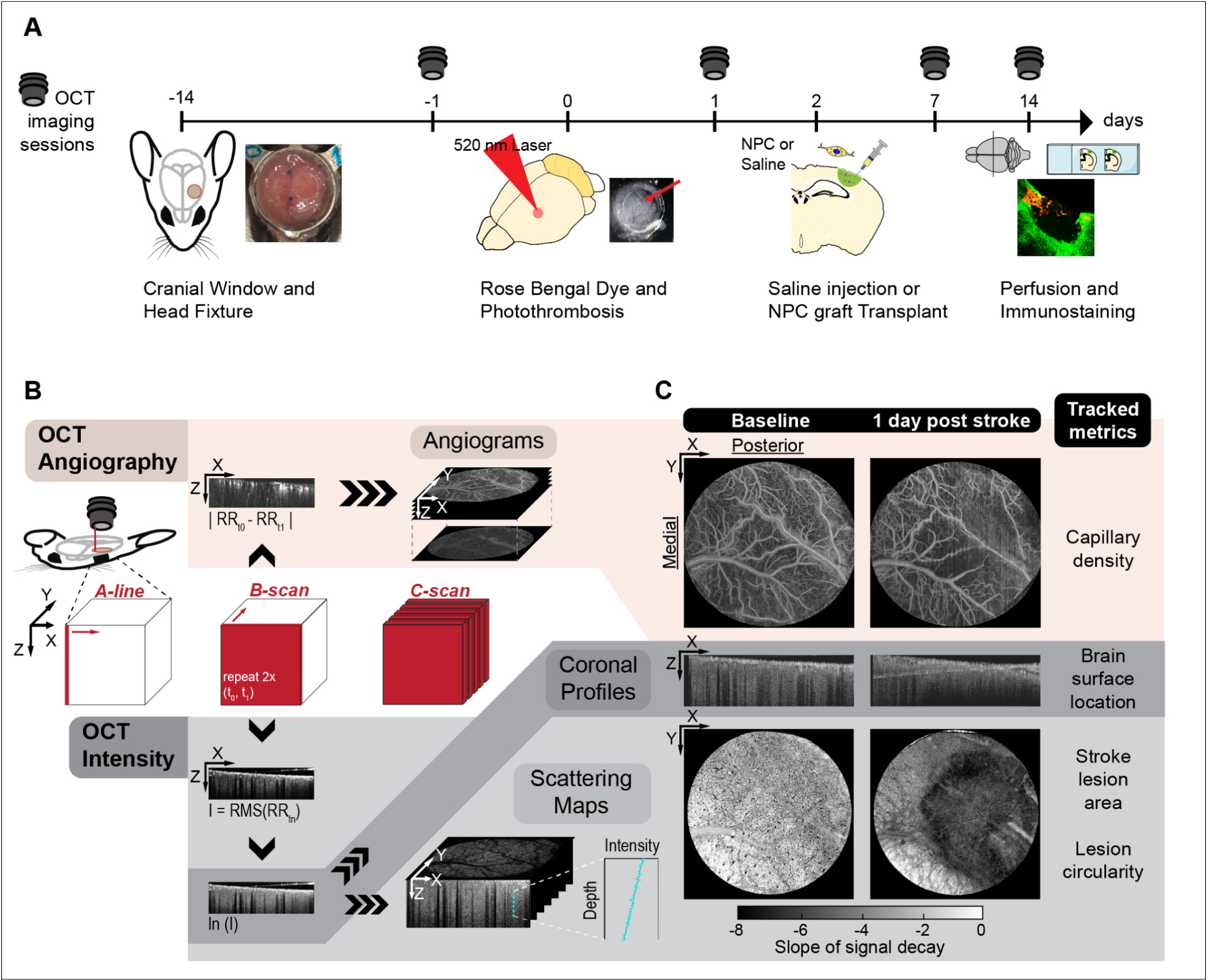
OCT experimental timeline and acquisition + processing steps. **A.** Experimental timeline. Initial window installation surgery is performed 2 weeks before photothrombotic stroke. Cell grafts or saline control are injected 2 days after stroke. Images are taken at baseline before stroke, then at 1, 7, and 14 days after stroke, at which time mice are perfused and tissue is processed for IHC. **B.** OCT acquisition and further processing of raw reflectivity (RR) data. Subtracting repeated B-scans isolates signal decorrelation from blood flow, which is then rastered into an angiogram. Alternatively, converting RR data to the natural logarithm of the signal intensity permits multiple analyses - (i) assessment of surface topography via coronal profiles, and (ii) linear fitting of the signal decay as a function of cortical depth, to assess optical scattering. **C.** Angiograms (top row), coronal profiles (middle row), and scattering maps (bottom row) before and after photothrombotic stroke. Right column: corresponding metrics extracted and tracked from the respective OCT images.

Firstly, OCT angiograms were generated by calculating the difference between two unique RR B-scans taken 20 milliseconds apart (RR_t0_ and RR_t1_), then rastering all results into a single C-scan (**Figure 1B**). The resultant OCT angiograms are images of functional, perfused vascular networks that are detected primarily based on the optical scattering properties of red blood cells (RBCs) flowing within the vasculature. Between temporally sequential B-scans (RR_t0_ and RR_t1_), OCT signal from perfused vasculature changes more dynamically than other neighboring tissue elements due to the mobility of RBCs. Subtracting RR_t0_ and RR_t1_ generates the OCT angiogram and results in high-contrast images of perfused healthy cortical vasculature in mice (Wang et al., 2007) (**Figure 1C**). Using a high numerical aperture (NA) objective lens (10x, NA = 0.28), we generated OCT angiograms that can resolve vascular networks down to the capillary level (2.5µm) when imaging healthy mouse cortex directly through the optical window. OCT angiograms generated from scans acquired at 24 hours after stroke have already been used to assess the acute severity of photothrombotic stroke lesions by tracking perfusion of capillaries adjacent to occluded pial vessels before and after stroke (Sunil et al., 2020). Here, using the same injury model and performing longitudinal imaging over the two weeks following stroke, we explored using OCT angiograms to also detect reperfusion in the stroke core and penumbra.

We also analyzed intensity (I) data from the acquired OCT scans, to extract a coronal profile within a single B-scan as well as OCT scattering maps from the entire image volume. Intensity (I) was defined as the root mean squared (RMS) of RR_t0_ and RR_t1_ acquired at each location. Coronal profiles were used to identify and quantify structural alterations to the tissue following stroke such as the separation of the meninges and brain surface from the imaging window (**Figure 1B**). In addition, OCT scattering maps, a measure of relative tissue scattering within different regions of the stroke, were extracted from the intensity data using the following procedure: The depth profile of this signal was initially parametrized as described in **Equation 1**, where μ*_b_* is the back-scattering coefficient, μ_*s*_ is the attenuation coefficient, *z* is depth, and ℎ(*z*) is the axial point-spread function (PSF). The ℎ(*z*) is a function of *z*, the focus depth *z*_*f*_, and the Rayleigh range *z*_*r*_, as shown in **Equation 2**. The μ_*s*_ is independent of the imaging system parameters and describes the attenuation of the OCT signal intensity as it travels through tissue or other media. As absorption of near-infrared light is negligible in mouse cortex, signal attenuation can be attributed solely to scattering, such that μ_*s*_ is equivalent to the scattering coefficient. Importantly, for OCT systems with sufficiently low numerical aperture, the confocal parameter, (i.e. twice the Rayleigh range), is larger than the maximum imaging depth. Thus, the ℎ(*z*) term can be neglected, and the equation for depth-dependent OCT signal decay can be simplified to a single exponential (Schmitt et al., 1993), as shown in **Equation 3**. Therefore, we exclusively analyzed images collected with a low-NA objective (5x, NA = 0.14) in this study to derive OCT scattering information, with images collected from 1000 x 1000 line scans over a 3 mm x 3 mm field of view. To minimize speckle noise, image volumes were downsampled in the axial (x-y) plane by a factor of 2 with bicubic interpolation, and 2 repeat volumetric scans were averaged together. Multiple additional assumptions were made to accommodate robust, fast, image processing. First, μ_*s*_ was assumed constant over an entire line scan. Also, local tissue scattering was determined by transforming the volume with a natural logarithm so that tissue scattering would vary linearly with depth instead of exponentially. Linear fits were then performed from 140 to 490 µm deep in the cortex, consistent with prior methods (Sunil et al., 2021), to avoid the fitting of unwanted scattering effects from the brain surface. Scattering was then reported as the slope of signal decay, with a steeper slope representing higher scattering. After mapping all decay coefficients across the imaging window, stroke lesion area was calculated by segmenting regions of higher scattering with an active contour algorithm (**SI** **Figure 1**). As an example, this procedure was used to map the emergent boundaries of a photothrombotic stroke lesion, 24 hours after stroke (**Figure 1C**). Here, we explored using OCT scattering maps to track changes in lesion size over multiple weeks as a metric of wound healing.

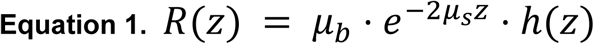

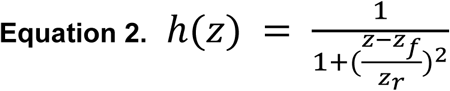

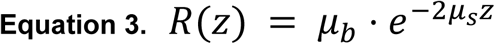

### 2. Longitudinal OCT imaging can track natural wound healing at stroke lesions

To evaluate how photothrombotic stroke lesions treated with saline evolve temporally during sub-acute wound healing, we generated and analyzed OCT angiograms (**Figure 2A,B**), coronal profiles through the lesion epicenter (**Figure 2C, SI Figure 1**), and OCT scattering maps (**Figure 2D, SI Figure 1**) at designated pre- and post-injury timepoints using the methodology described above. For the OCT angiograms, we processed two unique maximum intensity-projections in the axial (x-y) plane (MIPz) at depth ranges of 0 to 105μm (0-30 voxels) and 105 to 158μm (30-45 voxels) from the pial surface. The MIPz from 0 to 105μm (**Figure 2A**) typically reveals major pial vessels, minor meningeal vessels, and the emergence of intracortical capillaries in healthy neural tissue. Large pial vessels from the superficial MIPz were used as reference points for establishing the brain surface before and after stroke. Moving deeper, MIPz from 105μm to 158μm contained exclusively intracortical vessels (**Figure 2B**) and thus defined regions of neocortex whereby segmentation and evaluation of neurovascular capillary networks was possible. While the structural integrity and perfusion of major pial and minor meningeal vessels was preserved at all timepoints post-stroke in the surface MIPz, perfusion of intracortical capillaries was dramatically attenuated at 1 day after stroke, followed by a small and partial recovery of capillary density by 14 days. There were also notable alterations to the orientation of major pial vessels near the original stroke site over the 14 day evaluation period (**Figure 2A**). Angiograms of the deeper neocortical regions showed a similar loss of perfusion in intracortical capillaries after stroke revealing obvious stroke lesion margins at the 1 day timepoint (**Figure 2B**). To directly assess changes in capillary perfusion, we segmented capillary networks and computed network statistics including capillary network density (**Figure 2E.i**.) (Zudaire et al., 2011). As anticipated, loss of capillary perfusion after stroke was greater in the stroke core (as defined by OCT scattering maps) (Sunil et al., 2021) than in the surrounding tissue. At 1 day after stroke, there was a near total loss of capillary density (87.5%) in demarcated stroke lesion cores and a modest, but significant, 30.4% decrease in capillary density in the preserved neural tissue (**Figure 2E**). By 14 days post-stroke, there was essentially a complete return of capillary density in the preserved neural tissue region that matched baseline (healthy) neural tissue levels prior to stroke. However, the capillary density, as detected by the OCT angiogram, in the core region showed no significant recovery, maintaining an overall 70.1% decrease compared to healthy neural tissue baseline (**Figure 2E.ii**). Interestingly, unlike at 1 and 14 days, capillary density evaluations in processed OCT angiograms from scans at 7 days after stroke showed poor capillary segmentation outcomes and could not be readily analyzed. We attributed this poor segmentation to excessive signal loss due to scatter from the imaging path near the brain surface, yielding cortical OCT angiograms with a poor signal-to-noise ratio, which made the capillary signal less distinguishable from noise and motion artifacts. Despite these notable issues, these data show that OCT angiograms can be effectively used to track the spatial extent of impaired tissue perfusion in the different lesion compartments after stroke, as well as detect regions of the brain near strokes that become reperfused over the two-week, post-injury period.

**Figure 2:**
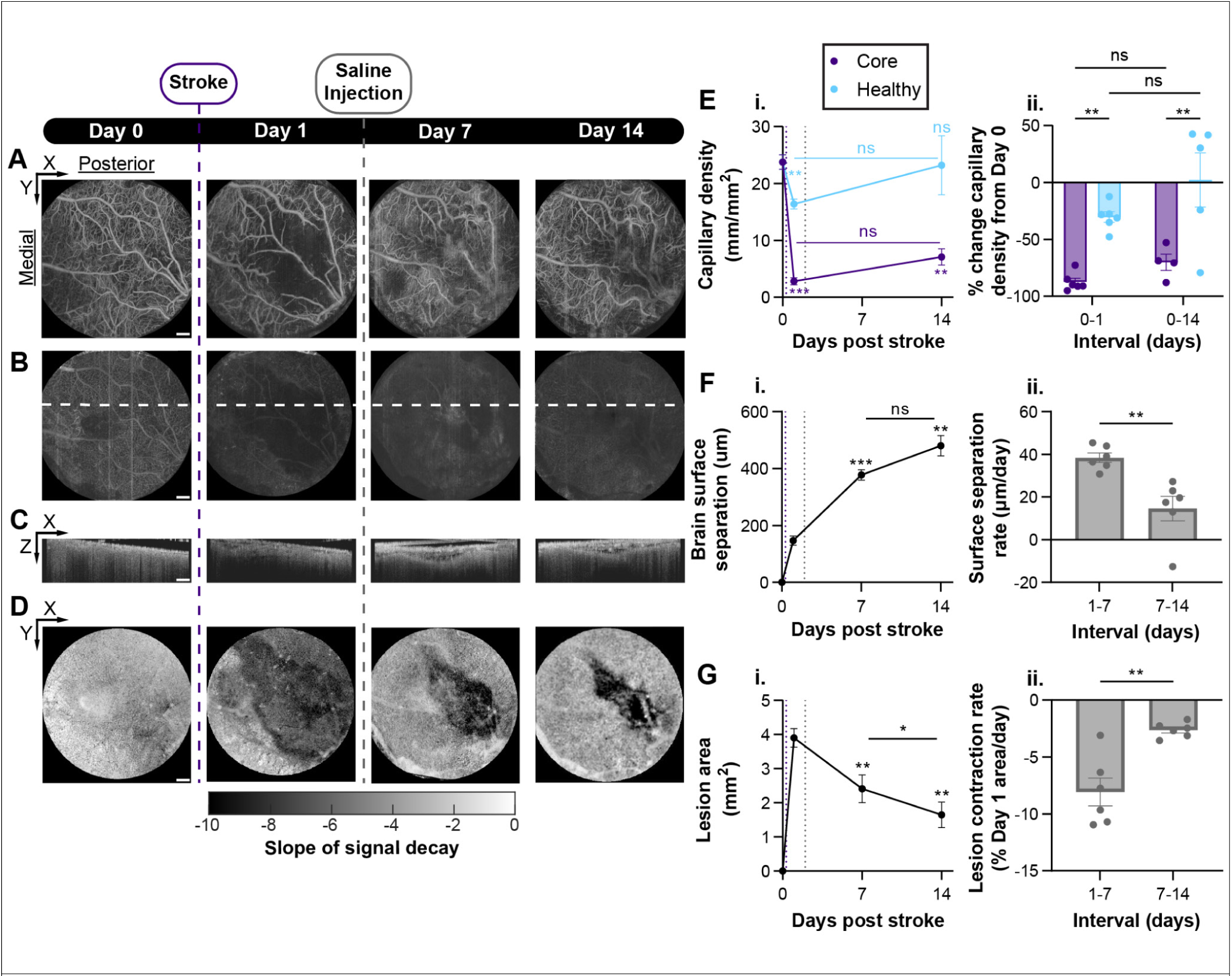
Representative longitudinal OCT imaging of photothrombotic stroke and tracked metrics of lesion evolution. **A.** Superficial maximum intensity z-projections (MIPz) (0-105 μm) of OCT angiograms. **B.** Intracortical MIPz (105-158 μm) of OCTA. Dotted line represents location of representative image slice shown in **C**. **C.** Representative Coronal profiles. **D.** OCT scattering maps generated from signal decay of OCT intensity. **E.** *i.* Longitudinal change in capillary density, (mm capillary/mm^2^ area), measured from intracortical OCTA MIPz acquired at 10x magnification, segmented as described, and subdivided into core/healthy ROIs according to scattering map-defined boundaries. Not significant (ns), *P < 0.034, **P < 0.01, ***P < 0.0001, mixed-effects analysis with Tukey multiple comparisons test. Line plot shows mean ± s.e.m., for N = 6 mice, or fewer if a given mouse did not have 10x images overlapping core or healthy tissue. *ii.* % Change in capillary density from baseline healthy measurements. Not significant (ns) and **P < 0.0021, mixed-effects analysis with uncorrected Fisher’s LSD. Bar chart shows mean ± s.e.m., with individual data points representing N = 6 mice, or fewer where indicated. **F.** *i.* Longitudinal change in maximum separation between brain surface and imaging window (μm), measured from coronal profiles sampled every 20 μm. Not significant (ns), **P = 0.0021, ***P < 0.0001, repeated measures one-way ANOVA with Tukey multiple comparison test, comparison to Day 1 or as otherwise indicated. Line plot shows mean ± s.e.m. for N = 6 mice. *ii.* Average surface separation rate during intervals between imaging timepoints. **P = 0.0065, paired t-test. Bar chart shows mean ± s.e.m., with individual data points representing N = 6 mice. **G.** *i.* Longitudinal change in stroke lesion area (mm^2^), measured from OCT scattering maps as described. *P = 0.024, **P < 0.0069, repeated measures one-way ANOVA with Tukey multiple comparison test, comparison to Day 1 or as otherwise indicated. Line plot shows mean ± s.e.m. for N = 6 mice. *ii.* Contraction rate of axial (x-y) stroke lesion area between imaging timepoints, relative to the lesion area initially measured 1 day after stroke. **P = 0.0087, paired t-test. Bar chart shows mean ± s.e.m., with individual data points representing N = 6 mice. All scale bars = 250 μm.

Next, we used coronal profiles, taken through the stroke lesion epicenter, to assess changes to the brain surface topography and the meningeal layers along the coronal (x-z) plane post-stroke. There were dramatic alterations to the brain surface that were isolated to the lesion environment but that changed dynamically across the different post-stroke evaluation time points (**Figure 2C**). Before stroke, the brain surface maintained complete and uniform contact with the imaging window, essentially creating a single visible interface. However, as early as 1 day after stroke, there was some but not complete separation of the brain surface from the imaging window such that two discrete interfaces were visible along various sections of the contacting surface. This separation of the brain surface from the window interface was hypothesized to be caused by localized swelling and inflammation of the meningeal tissue layer and fluid extravasation leading to edema, since pockets of fluid could be identified above the brain surface as regions showing near-zero OCT intensity, which is consistent with how fluid filled spaces appear on OCT. The meninges were identified as the first visible interface beneath the optical window, while the brain surface was identified as the deepest visible interface detected on the coronal profiles, beneath which steady signal decay indicative of cortex was observed. The tissue separation induced by injury was exacerbated by 7 days after stroke and beyond, such that the stroked tissue region could be readily identified as a notable divot in the cortex by OCT, with this region of the brain surface being maximally contracted away from the optical window. Above the vacated divot at the stroke core at 7 and 14 days, there were multiple, separate, highly reflective interfaces. This likely arose from exacerbated inflammation within the meningeal layers and widespread fluid extravasation. Contraction of the brain surface at the stroke was associated with the tissue separation, and measured as the maximum distance between the imaging window and the brain surface, was tracked and quantified temporally. While at 1 day the average separation of the brain surface from the imaging window was 172 µm, by 7 days this separation had more than doubled to reach 406µm (**Figure 2F.i.**). There was continued increased separation at the lesion over the course of the subsequent week, resulting in total average separation of 534µm, but the rate of brain surface contraction had significantly slowed over the second week compared to the 1 to 7 days post-stroke interval, by a factor of 2.5 (**Figure 2F.ii.**).

The alterations to the brain surface topography at the lesion during the sub-acute wound repair period required us to adjust the image processing methodology used to compute the OCT scattering maps. For images taken at healthy baseline, the brain surface, flush with the window, could be readily fit and represented by a plane, which permitted the necessary correction for any tilt of the brain surface relative to the microscope objective that is inherent to the imaging set-up. By contrast, at sub-acute post-stroke timepoints, divot formation and tissue separation at the stroke lesion resulted in brain surfaces that had complex, undulating topographies. To account for these brain surface perturbations, for each C-scan we manually applied surface contours on serially sampled coronal (x-z) B-scans and performed bicubic interpolation to generate a smooth surface mesh. Computing the cortical depth relative to this surface mesh, as opposed to a flat plane, significantly improved the quality of processed OCT images in multiple ways. Firstly, it facilitated accurate, corroboratory mapping of distinct layers of cortical vasculature with OCT angiograms, by accounting for structural perturbations to the cortex (**SI** **Figure 2**). Secondly, OCT scattering maps could be acquired as previously described, but with the cortical depth range now relative to the surface mesh instead of the imaging window. This eliminated erroneous inclusion of meningeal tissue in the scattering analysis, so that cortical tissue could be analyzed in isolation as in the uninjured state. As a result, contrast between healthy and ischemic tissue was improved. Together, these data show that coronal profiles can be used to track interfaces such as the meninges and brain surface, in the event they separate from the imaging window or each other following stroke. Our results demonstrate that tracking the spatial evolution of the brain surface is crucial to both improving OCT image processing and assessing the extent of meningeal inflammation and cortical contraction post stroke.

To measure stroke lesion size and any extent of infarct contraction in the axial (x-y) plane over time, we generated OCT scattering maps as described above, from 140µm beneath the brain surface to 490µm, or at the point at which signal linearity was lost, if it was shallower. Then, we plotted each calculated slope of the linear fit as a pixel value in the axial (x-y) plane, and segmented regions of steep decay (i.e. high scattering) to define the boundary between stroke lesion core and preserved neural tissue. This processing approach enabled assessment of the longitudinal changes in lesion area within this plane to be readily identified (**Figure 2D**). Using this approach, we noted that the lesion area was greatest at 1 day after stroke and that there was persistent contraction of the lesion in the axial plane over the 14-day evaluation period (**Figure 2G.i.**). Notably, the lesion contraction rate was 3 times faster between 1 and 7 days after stroke compared to the following week (**Figure 2G.ii.**). These data demonstrate the utility of OCT scattering maps for tracking lesion size and shape in the axial plane and document the significant contraction of untreated stroke lesions that naturally occurs during the sub-acute wound repair phase.

### 3. NPC grafts alter the wound repair of photothrombotic stroke lesion cores

Previously, we identified that acute grafting of neural progenitor cells (NPC) into mouse striatal stroke lesions made using L-NIO injections directs enhanced localized glial repair that restricts inflammation and fibrosis at non-neural lesion cores (Adewumi et al., 2024; O’Shea et al., 2022). We hypothesized that such NPC graft-induced alterations to the stroke lesion environment may be detectable by OCT, and if so, would enable us to perform longitudinal monitoring of graft-induced changes. However, before assessing whether OCT is capable of non-invasively tracking NPC-induced changes, we first sought to verify that similar NPC-derived glia repair outcomes as seen in L-NIO strokes are conferred when grafts are made into acute photothrombotic stroke lesions, since we have established that grafting outcomes can be dramatically affected by different types of lesion environments (Adewumi et al., 2024). To do this, we grafted NPC into photothrombotic strokes at two days after inducing the injury and compared the resultant lesion phenotype against saline treated controls by immunohistochemistry at two weeks post-stroke (**Figure 3A-D**). As in our prior studies, we transplanted NPC derived from a RiboTag transgenic mouse embryonic stem cell line such that all grafted cells and their progeny constitutively express a hemagglutinin tag (HA). Detection of graft-derived HA-expression by IHC enabled post-hoc analysis of graft survival, anatomical location, and contributions to wound repair.

**Figure 3:**
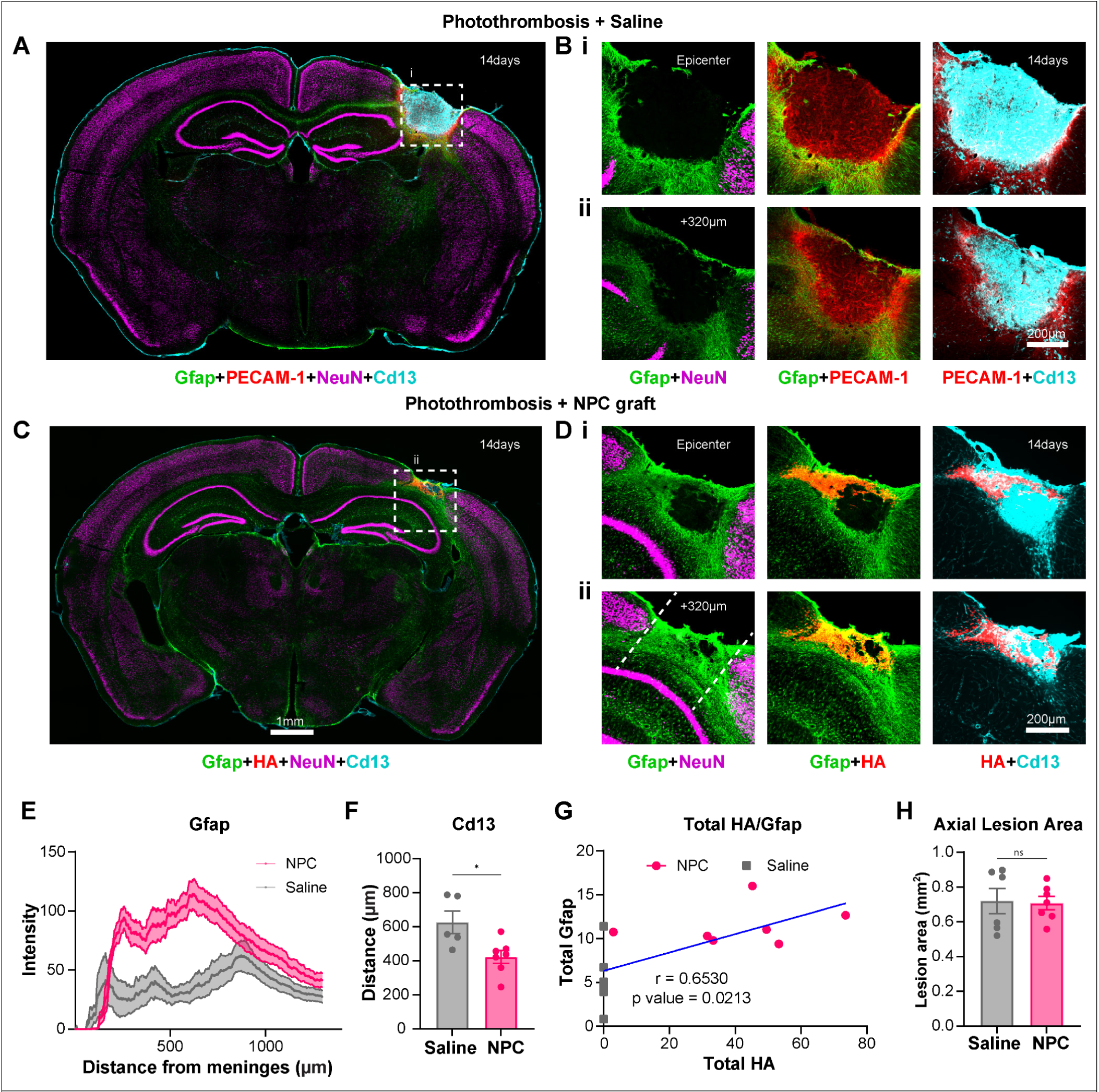
Immunohistochemistry of stroke lesions and spatial analysis of lesion phenotype alteration by NPC grafts. **A**. overview image of coronal brain section for NPC grafts at 2 weeks. **B**. Zoomed in image of grafted lesions (**i**) epicenter of lesion (**ii**) 320 µm from epicenter of lesion (inset from A); showing stains for Gfap (astrocytes), HA (grafted NPCs), NeuN (neurons) and Cd13 (macrophages). **C**. Overview of brain section for Saline group at 2 weeks. **D**. Zoomed in images of lesion (**i**) Epicenter of lesion (inset from C) (**ii**) 320 µm from epicenter of lesion; showing stains for Gfap (astrocytes), PECAM-1 (vasculature), NeuN (neurons) and and Cd13 (macrophages). **E**. Gfap trace of NPC and saline groups radially from meninges into the cortex. **F**. Half width maximum of Cd13 trace (Student’s t test; p value-0.0182). **G**. Correlation graph of total HA (Cell grafts) and total Gfap (r value: 0.6530, p value: 0.0213). **H**. Axial lesion area derived by neuron-devoid tissue in X-plane. Sample size n=6 for saline, n=7 for NPC. Student’s t-test, ns=not significant

By two weeks post injury induction, saline treated photothrombotic stroke lesions exhibited focal neuron and glial cell loss within a defined region in the cortex that extended from the brain surface through the full thickness of the cortical column resulting in a lesion that was approximately 1000µm deep (**Figure 3A**). Regions of tissue damage were remodeled into compartmentalized lesions comprising a dense core of Cd13-positive macrophage and fibroblasts that were separated from preserved neuron-containing neocortex by a thin astrocyte border. For these two-week post injury lesions, dense PECAM-1-positive angiogenic vessels penetrated into and throughout the non-neural lesion compartment but these newly formed vessels were thin, tortuous, and lacked glia-limitans structures which made them distinctly different to characteristic neurovasculature that was apparent in the preserved neural tissue adjacent to stroke lesions (**Figure 3B**). NPC grafted into the same photothrombotic strokes survived in moderate numbers and were retained locally within the lesion core at two weeks but were mostly identified in superficial cortical regions such that essentially all HA-positive grafted cells and their progeny were restrained to within 500µm of the brain surface (**Figure 3D, SI Figure 3F**). By two weeks post-stroke, all NPC had autonomously differentiated into Gfap-positive wound repair astrocytes that contributed to an increased total number of Gfap-positive cells within the lesion (**Figure 3E, SI Figure 3C**), which was a response that positively correlated with total HA-positive grafted cell survival (**Figure 3G**). Graft-derived astrocytes modified the non-neural lesion core environment in a manner that was consistent with, but somewhat less effective than, what we observed in our previous studies using L-NIO striatal strokes (**Figure 3C&D**) (Adewumi et al., 2024; O’Shea et al., 2022). This conferred effect resulted in an anisotropic contraction of the lesion when viewed via histology, with NPC grafts resulting in lesions that were more contracted along the z axis (perpendicular to the imaging window) (**SI Figure 4**). However, total graft survival at photothrombotic strokes was far greater than the poor survival we have observed previously within collagenase-induced hemorrhagic stroke lesions in the striatum at the same post-injury time point, suggesting that photothrombotic strokes, which result in severe ischemic infarcts, are overall less hostile towards grafts compared to severe hemorrhagic lesions (**SI Figure 5**). NPC treatment of photothrombotic strokes resulted in a modest but significant reduction in Cd13-positive inflammation and fibrosis at the lesion compared to the saline treated group, leading to overall smaller non-neural lesion cores. (**Figure 3F**, **SI Figure 3A,B**). For both groups of mice, there was significant ipsilesional cortical volume loss due to tissue atrophy at, and immediately surrounding, the stroke lesion which caused an inward deviation of brain surface in this region. Within the core region, NPC graft-treated lesions were notably contracted along the dorsal-ventral axis compared to saline-treated strokes. However, in the axial plane, there was no significant difference in the width of the neuron-devoid lesion tissue between the saline and NPC treated groups suggesting that grafted cells did not directly repopulate lesions with neurons or increase the survival of host neurons when transplanted into strokes two days after injury, which was consistent with our previous observations (**Figure 3H**).

Overall, the IHC-based analysis at two weeks post-injury showed that NPC graft treatment, when injected into acute photothrombotic strokes at two days after stroke onset, contributed to the overall astroglial wound response at stroke lesions which helped to restrict the size of the resultant non–neural lesion core. Based on these observations, we were motivated to test whether we could use OCT imaging to longitudinally track these NPC graft-induced effects at stroke lesions.

### 4. Longitudinal OCT imaging detects NPC graft-induced changes at stroke lesions

To longitudinally evaluate NPC treatment effects using OCT we again transplanted cell grafts two days after the induction of stroke. Stroke lesions receiving NPC grafts or saline were of comparable size, as assessed by OCT scattering maps, at one day post-stroke suggesting that any effects that could be seen in the NPC treatment group were unlikely to be due to any significant differences in the conferred injury between the two groups (**Figure 4G.i.**). As before, we longitudinally tracked the effects of NPC treatment across the sub-acute wound healing period by generating OCT angiograms to evaluate capillary loss and reperfusion at the core and adjacent preserved neural tissue (**Figure 4A**,**B**), coronal profiles through the lesion epicenter to assess brain and meningeal surface changes (**Figure 4C**), and OCT scattering maps to analyze stroke lesion size changes in the axial plane (**Figure 4D**).

**Figure 4:**
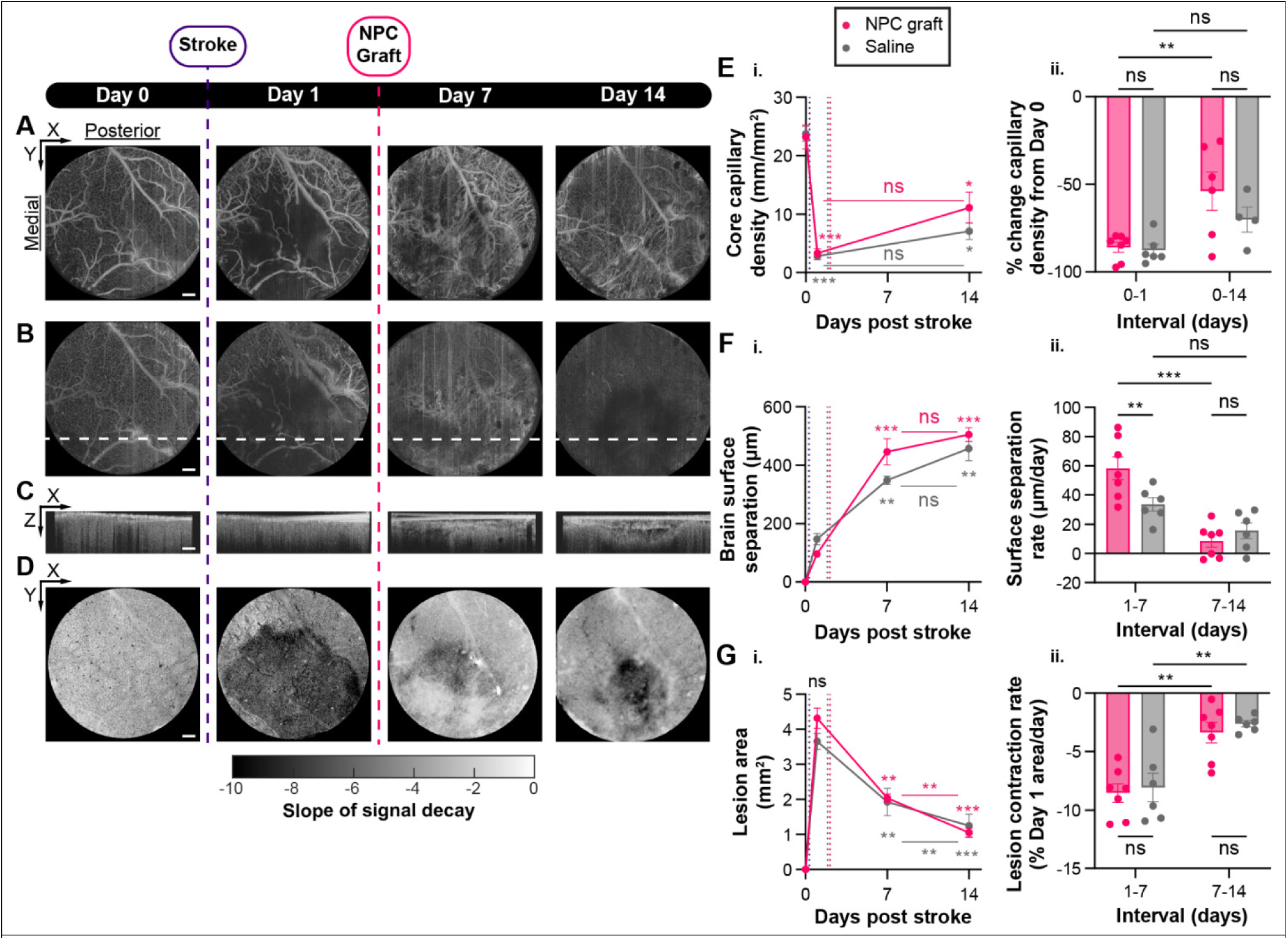
Representative longitudinal OCT imaging of photothrombotic stroke before and after NPC graft treatment, and measurements of altered lesion evolution. **A.** Superficial maximum intensity z-projections (MIPz) (0-105 μm) of OCTA in an NPC graft recipient. **B.** Intracortical MIPz (105-158 μm) from the same OCTA scan. Dotted line represents location of representative image slice shown in **C**. **C.** Representative Coronal profiles. **D.** OCT scattering maps generated from signal decay of OCT intensity. E. *i.* Longitudinal change in capillary density in the stroke core, (mm capillary/mm^2^ area), measured from intracortical OCTA MIPz acquired at 10x magnification and segmented as described. Not significant (ns), *P < 0.015, ***P < 0.0002, mixed-effects analysis with Sidak’s multiple comparisons test. Colored comparisons are to said group’s Day 1 or as otherwise indicated. Line plot shows mean ± s.e.m., for N = 7 NPC graft mice and N = 6 saline mice, or fewer if a given mouse did not have 10x images overlapping core tissue. *ii.* % Change in capillary density from baseline healthy measurements. Not significant (ns) and **P < 0.01, mixed-effects analysis with uncorrected Fisher’s LSD. Bar chart shows mean ± s.e.m., with individual data points representing N = 7 graft mice or N = 6 saline mice, or fewer where indicated. F. *i.* Longitudinal change in maximum separation between brain surface and imaging window (μm), measured from coronal profiles sampled every 20 μm. Not significant (ns), *P < 0.048, **P < 0.0059, ***P < 0.0007, repeated measures two-way ANOVA with Tukey multiple comparison test. Comparisons in black are between groups at a single time point. Colored comparisons are to each group’s Day 1 or as otherwise indicated. Line plot shows mean ± s.e.m., for N = 7 NPC graft mice and N = 6 saline mice. *ii.* Average surface separation rate during intervals between imaging timepoints. Not significant (ns), **P < 0.0068, ***P < 0.0003, repeated measures two-way ANOVA with Tukey multiple comparison test. Bar chart shows mean ± s.e.m., with individual data points representing N = 7 graft mice or N = 6 saline mice. G. *i.* Longitudinal change in stroke lesion area (mm^2^), measured from OCT scattering maps as described. Not significant (ns), *P < 0.0132, **P < 0.0033, ***P < 0.0005, repeated measures two-way ANOVA with Tukey multiple comparison test. Comparisons in black are between groups at a single time point. Colored comparisons are to said group’s Day 1 or as otherwise indicated. Line plot shows mean ± s.e.m., for N = 7 NPC graft mice and N = 6 saline mice. *ii.* Contraction rate of axial (x-y) stroke lesion area between imaging timepoints, relative to the lesion area initially measured 1 day after stroke. Not significant (ns), **P < 0.005, repeated measures two-way ANOVA with Tukey multiple comparison test. Bar chart shows mean ± s.e.m., with individual data points representing N = 7 graft mice or N = 6 saline mice. All scale bars = 250 μm.

The two unique MIPz OCT angiograms showing vascular structures at the meninges and brain surface (**Figure 4A**) as well as neocortex (**Figure 4B**) respectively were not significantly altered by NPC treatment compared to the saline control. As before, capillary density was dramatically attenuated at one day after stroke in the core region, which was followed by a small but insignificant increase in capillary density due to NPC treatment by 14 days (**Figure 4E**). The transient attenuation in capillary density in preserved neural tissue that was seen for NPC treatment was also observed with saline treatment (**SI** **Figure 6**). Interestingly, staining the tissue at two weeks for the endothelial maker, PECAM-1, using post hoc IHC revealed that for both groups there was a dense network of remodeling capillaries within the lesion core environment that was not readily detected by OCT angiography.

Vasculature density by PECAM-1 staining was not noticeably altered by NPC treatment. While there were numerous vessel-like structures detected within lesion environments, including many that were interacting with HA and Gfap-positive graft derived astrocytes, these angiogenic vessels lacked the morphology, organization and density of healthy neurovasculature (**SI Figure 7**). These structural differences, plus the potential for altered RBC flow dynamics in these angiogenic vessels, could be responsible for the failure to detect them on OCT angiograms. It is also possible that reductions in SNR due to optical scattering affect our ability to properly resolve OCT angiograms within stroke cores.

These data broadly suggested that, at least by OCT detection, NPC grafts failed to rescue vascular function in ischemic stroke cores in this photothrombotic stroke model.

In contrast to the OCT angiography, evaluation of OCT-derived coronal profiles demonstrated significant NPC graft-induced changes to the brain surface topography and the meningeal layers compared to saline treatment (**Figure 4F**). One day after stroke and prior to NPC treatment, the extent of separation of the meninges and brain surface from the window interface was no different between the two groups. However, in the time period between 1- and 7-days post-stroke (5 days after NPC graft treatment) we detected a significantly greater rate of brain separation in the NPC graft-treated mice that was 74% greater than the saline controls over the same interval. This same trend was observed for the meninges, with a 106% greater separation rate in the NPC graft-treated mice. However, tissue separation differences between NPC graft and the saline control were not significantly different by two weeks (**SI Figure 8**). The cortical contraction induced by NPC grafts was most pronounced during the first week after treatment, with a mean tissue separation from the window of 446 µm, and there was only a modest 60 µm further increase in tissue separation over the course of the subsequent week that was less than the 109 µm observed in the saline group during the second week. As a result, the ultimate difference in cortex-window separation between the two groups after two weeks was reduced to 10%. The IHC evaluations reported above noted an increased lesion contraction of approximately 200µm in this plane for NPC graft treatment, suggesting a possible relationship between tissue separation from the optical window and mechanical contraction of the lesion.

Despite notably increased contraction at the lesion epicenter in the coronal plane in NPC graft-treated mice, quantification of the total lesion area in the axial plane revealed by the OCT scattering maps showed no significant differences between the groups at any of the imaging timepoints across the 2 week post-injury period (**Figure 4G.i.**). Additionally, the contraction rates calculated between each of the imaging timepoints were not significantly different for NPC graft or saline control (**Figure 4G.ii.**).

Longitudinally acquired scattering maps were assessed for possible presence of anisotropic contraction in the axial (x-y) plane due to grafting. However, we found no significant difference between groups in the changes in lesion circularity over time, suggesting that the NPC treatment did not substantially distort the natural temporal pattern of lesion shrinkage (**SI Figure 9**).

These data show that NPC graft-induced changes to stroke lesions can be detected by some, but not all, OCT parameters and that key differences between NPC and saline groups are consistent with features observed in post-hoc IHC analysis. Surprisingly, despite notable angiogenesis in stroke lesion cores, OCT angiography was unable to identify these vessels at any of the post-injury time points evaluated.

### 5. Key stroke lesion parameters derived by OCT correlate minimally with IHC readouts

Up until now, post hoc IHC has been the gold standard in the neurobiology community for evaluating the size of strokes and the cellular phenotypes within the multi-cellular compartments that make up the stroke lesion. The ability to use intravital imaging like OCT to derive similar information about strokes would enable longitudinal study of lesion remodeling in an animal-specific manner while reducing the reliance on multiple histological endpoints. However, to achieve this requires establishing the specific cellular information that OCT imaging parameters can provide at stroke lesions.

Additionally, a notable limitation of the mouse photothrombotic stroke model is that the extent of the conferred tissue damage is highly dependent on the morphology and coverage of the vessel targeted for selective occlusion (Sunil et al., 2020). As a result, there can be significant inter-animal variation in stroke lesion injury severity within a given group of mice that can hamper the ability to detect modest treatment effects by pairwise comparison using standard pre-clinical sample sizes. While such a large variance in lesion severity is not ideal for evaluating treatment effects and may have impacted our ability to detect NPC-dependent outcomes by OCT in this current study, it opens up the possibility to perform a robust correlation analysis between OCT and IHC parameters from these mice due to the extended range of lesion sizes. Thus, we next compared OCT imaging results with IHC imaging results for each animal at the common two-week time point to assess whether we could correlate any specific metrics from each of the two evaluation methods and identify key cellular features at lesions detected by OCT.

Preliminary inspection of small and large lesions from saline control mice showed notable concurrence in the general size of the lesions detected by both OCT and the multicellular IHC at two weeks, with large lesions determined by OCT (e.g. mouse S3) also appearing large by IHC even though OCT scattering maps resolve an axial view (optical x-y plane) of the lesion while IHC sections provide a coronal assessment (optical x-z plane) (**Figure 5A**). However, for NPC treated mice, the largest lesion determined by OCT was not detected as such by IHC (e.g. mouse N4), while one of the smallest lesions determined by OCT (mouse N6) was assessed to be the largest by IHC (**Figure 5B**). To explore the extent to which there was any correlation between OCT and IHC readouts more thoroughly at two weeks, we separated the IHC-based lesion characterizations into three distinct parameters and compared their correlation with two unique OCT-derived parameters that showed a robust dynamic range across the cohort. For the IHC readouts, we used: (i) IHC lesion area which was derived by a stereological analysis of the NeuN-negative lesion regions across three coronal sections spaced 320µm apart; (ii) the total Gfap-positive tissue detected at the lesion; and (iii) the total size of the Cd13-positive non-neural core within the lesion. We explored a pair-wise correlation analysis between each of these IHC parameters and (i) OCT scattering maps defining lesion size in the axial plane (**Figure 5C**), or (ii) brain surface separation revealing lesion contraction from OCT coronal profiles (**Figure 5D**). For these pair-wise correlation analyses, we pooled NPC treated and saline control mice together to increase the sample size and the power of the correlation comparison since all readouts employed here profiled features of the lesion common to both groups and nothing specific to the graft treatment itself. We anticipated that the IHC lesion area parameter, which provided the most accurate IHC-based estimate of lesion size within the axial plane, would be the IHC parameter most likely to correlate with the OCT lesion area results, since both readouts provided information on the lesion size in the axial plane. However, while there was modest, positive correlation between these two lesion area parameters (r >0.5), it was not statistically significant (p value, 0.0772). Likewise, total Gfap at the lesion showed no correlation with the lesion area measured by OCT despite there being large and significant differences in this parameter between NPC graft and saline control (**Figure 5C**). By contrast, the total size of the Cd13-positive, non-neural lesion core showed significant, positive correlation (r∼0.78, p value, 0.0028) with the OCT lesion area parameter (**Figure 5C**). Likewise, the OCT-derived surface separation data also only showed significant correlation with total Cd13 and not with the other two IHC-based parameters (**Figure 5D**). However, while the lesion size showed positive correlation with total Cd13, the surface separation was negatively correlated with total Cd13 such that smaller lesions that had reduced total Cd13 showed increased total surface separation.

**Figure 5:**
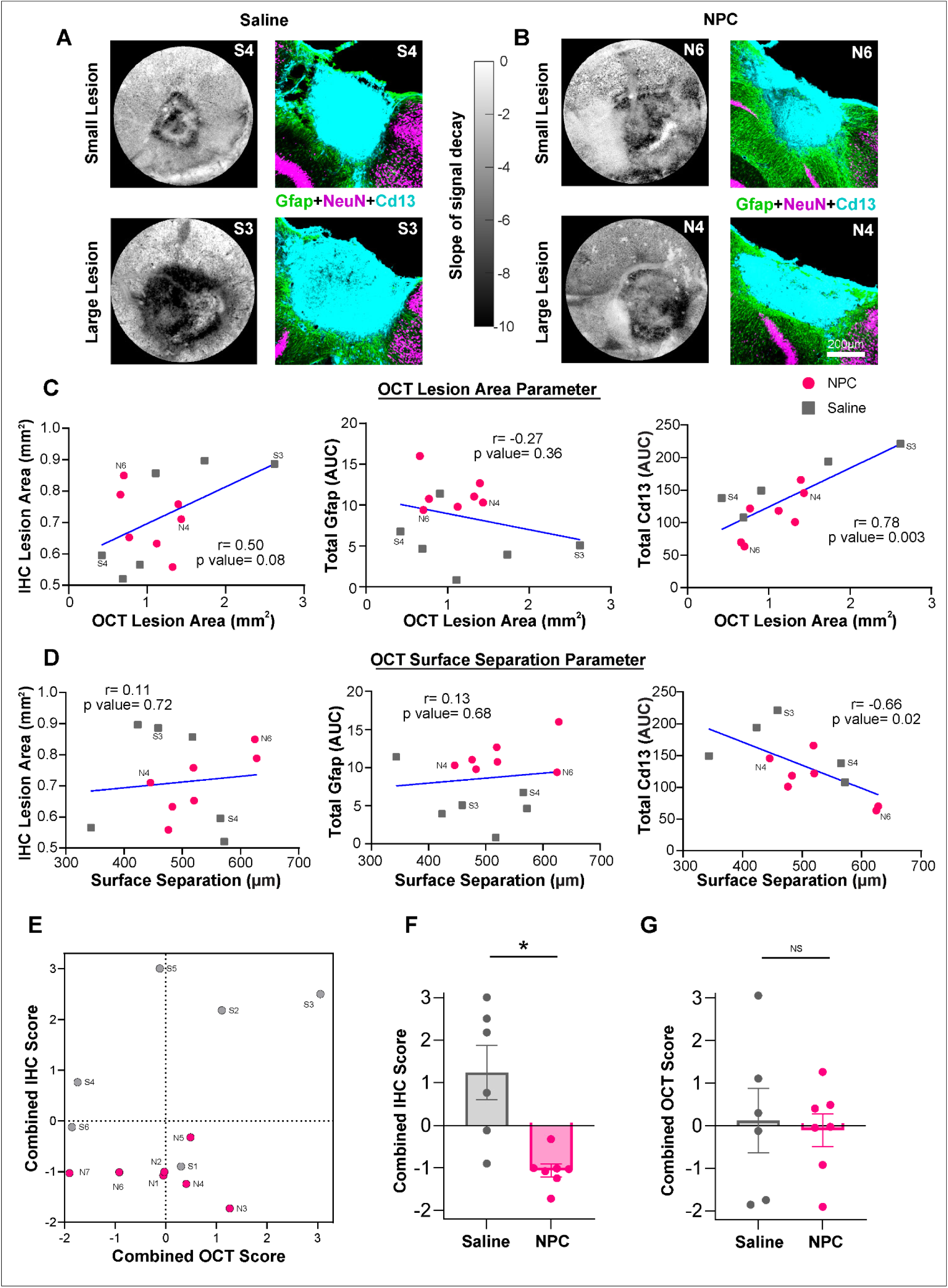
Correlation between OCT and IHC measurements at 2 weeks. **A.** OCT scattering maps and Immunohistochemistry profiles for smallest and largest lesions in Saline group at 2 weeks. **B.** OCT scattering maps and Immunohistochemistry profiles for smallest and largest lesions in NPC graft group at 2 weeks. **C.** Correlation scatter plot of OCT lesion area parameter and **(i)** IHC lesion area **(ii)** Total Gfap by IHC **(iii)** Total Cd13 by IHC. **D.** Correlation scatter plot of OCT surface separation parameter and **(i)** IHC lesion area **(ii)** Total Gfap by IHC **(iii)** Total Cd13 by IHC. **E.** Comparison of Combined IHC and OCT PCA scores for saline control and NPC graft treated mice. A larger positive number for both metrics indicates a more severe lesion. **F.** Bar graph comparing the mean combined IHC score for saline control and NPC graft treatment. *P = 0.0145, Welch’s t-test. **G.** Bar graph comparing the mean combined OCT score for saline control and NPC graft treatment. Not significant (ns), Welch’s t-test.

Given the multicellular nature of CNS injuries like stroke, it is likely that lesion severity may not be adequately described by any one single parameter from either OCT imaging or IHC analysis. To determine if a combination of parameters could improve the extent of correlation between the OCT and IHC readouts, we next incorporated multiple factors into a single combined score for OCT and IHC separately using principal component analysis (PCA). To perform this analysis, for each individual mouse in the study we incorporated data from three unique OCT parameters (OCT lesion size, OCT surface separation, OCT circularity) and four unique IHC parameters (IHC axial lesion area, total Gfap, total Cd13, and coronal lesion depth). Performing PCA on the three OCT parameters resulted in a first principal component (PC1) that described 60.4% of the total system variation while the PCA applied to the four IHC-based parameters resulted in a PC1 that described 58.32% of the total system variation, both of which we considered adequate for incorporating the multiple different parameters into a single evaluation metric (**SI Figure 10**). Factor scores along PC1 were used to define each of the OCT and IHC combined scores for each animal, with a larger positive number indicating a larger, more severe lesion (**Figure 5E**). While both scores identified the largest lesion (sample S3) there was no overall correlation between these two combined scores across the entire cohort of mice (**Figure 5E**). Notably, analyzing the distribution of the combined scores for NPC graft and saline controls separately revealed that the combined IHC scores were significantly different between the two groups (**Figure 5F**) but there was essentially no difference between the two groups on the combined OCT score (**Figure 5G**).

These data show that while total Cd13 by IHC had a significant positive correlation with two unique OCT derived parameters when assessed at the same two-week post-stroke timepoint, we identified limited overall correlation between the two evaluation methods across multiple individual and combined metrics. Furthermore, while NPC graft treatment effects could be detected by a combined assessment of multiple IHC parameters, OCT failed to identify these differences, suggesting that there may be key features of treatment-induced lesion remodeling that are not readily detectable by OCT using the current imaging processing methodology.

### 6. Acute post-stroke OCT parameters are prognostic of NPC grafting outcome

Non-invasive OCT imaging could become an important preclinical or clinical tool for guiding decision making on therapeutic interventions for stroke including cell grafting treatments. Two scenarios where OCT would be useful for guiding decision making about new treatments include: (i) generating prognostic information about the lesion environment to aid assessments of the suitability of that lesion for a particular treatment at that particular time, and (ii) for tracking whether a treatment is having an effect early after administration. For each mouse receiving the NPC graft treatment two days after stroke, we detected pronounced and consistent contraction of lesions over the two week post-stroke time-course, as measured by OCT scattering maps. By OCT, we detected regions of abnormally high or low scattering adjacent to the lesion at 14 days after stroke in some animals and were likely due to limitations in our OCT scattering model, which assumes that scattering is constant with depth in the tissue and may fail to fully recapitulate the cellular heterogeneity of the subacute stroke lesion environment. Additionally, across the NPC graft-treated cohort there was considerable variation in the number of HA-positive, graft-derived cells detected by IHC (**Figure 6A**). In our previous studies, we noted that graft survival is significantly poorer in severe versus moderate ischemic lesions (Adewumi et al., 2024), raising the prospect that lesion size directly affects grafting outcomes. To test if NPC survival noted at 2 weeks could be predicted by using acute OCT scattering measurements to characterize the lesion, we performed a correlation analysis between the pre-treatment (Day 1) lesion area derived by OCT scattering measurements and total HA-positive graft-derived cell survival detected by IHC at two weeks. Interestingly, there was significant inverse correlation between these parameters such that larger lesions by OCT resulted in lower total numbers of surviving HA-positive graft derived cells (**Figure 6B**). These data are consistent with, and further reinforce, our previous observations that lesion size and severity are powerful determinants of grafting outcome in stroke. Practically this also shows that OCT scattering measurements taken prior to treatment could be used to predict cell graft survival simply by computing the overall size of the acute lesion area.

**Figure 6:**
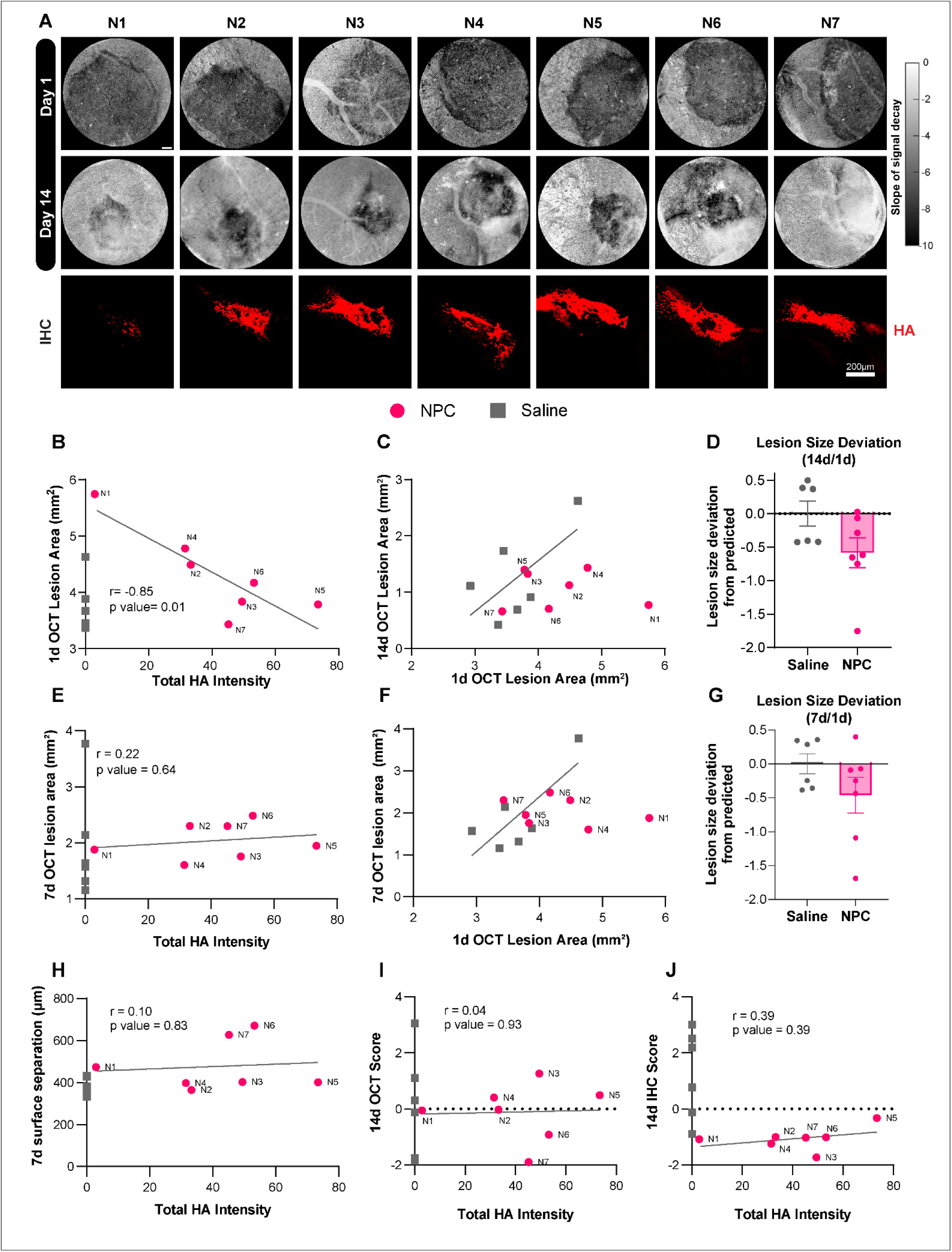
Prognostic potential of OCT for cell grafting outcomes. **A.** Top: OCT scattering maps generated as previously described for all NPC graft recipients, at 1 day and 14 days after stroke. Scale bar = 250 μm. Bottom: corresponding HA stain. **B.** Correlation scatter plot of total HA and lesion area measured by OCT 1 day after stroke. (r value: -0.8524, p value: 0.0148) **C.** Correlation scatter plot of lesion area measured by OCT 1 day and 14 days after stroke. Line of best fit is for saline group only. **D.** Deviation of Day 14 stroke lesion sizes from prediction, determined by vector distance from the best fit line. **E.** Correlation scatter plot of total HA and lesion area measured by OCT 7 days after stroke. (r value:0.2196, p value: 0.0148) **F.** Correlation scatter plot of lesion area measured by OCT 1 day and 7 days after stroke. Line of best fit is for saline group only. **G.** Deviation of Day 7 stroke lesion sizes from prediction, determined by vector distance from the best fit line. **H.** Correlation scatter plot of total HA and surface separation measured by OCT 7 days after stroke. (r value: 0.1030, p value: 0.8261) **I.** Correlation scatter plot of total HA and OCT score from PCA. (r value: 0.0424, p value: 0.926) **J.** Correlation scatter plot of total HA and IHC score from PCA. (r value = 0.385, p value = 0.393)

Given the prognostic power of the 1d OCT lesion measurements for overall cell graft survival, we next wanted to test whether this acute lesion measurement could also be used to predict other outcomes such as how individual lesions change over time. Plotting 1d OCT measurements against lesion size measurements at either 7 or 14 days revealed the extent to which acute lesion size influenced the lesion phenotype at these later time points both with and without the graft treatment intervention. Applying a linear fit to the saline control mice on these plots could serve as a predictive line of natural lesion area contraction during the subacute wound healing phase (**Figure 6C**). NPC treated stroke lesions showed a notable deviation from this natural trajectory at 14 days with all but one NPC treated mouse (i.e. 6/7 mice) falling below the predictive line such that the 14d lesions were smaller in size than would otherwise be predicted if the NPC graft treatment was not administered (**Figure 6D**). However, despite a notable trend towards lesion reduction from NPC treatment this was not statistically significant, likely due to the considerable variability among the contraction ratios of the saline controls. The complicated synergy between cell graft efficacy and stroke lesion environment is emphasized here-NPC grafts have the capability to augment the natural wound healing response to stroke, but are ultimately affected themselves by the size and severity of the stroke lesion. The demonstrated relevance of acute stroke lesion characteristics suggests a vital role for intravital imaging tools such as OCT, which can reliably and non-invasively measure lesion size, shape, and location in the acute phase.

The other potential use of longitudinal OCT for evaluating NPC graft outcomes is in the monitoring of their conferred effects throughout the subacute wound healing phase. To assess the viability of OCT imaging for this application, we next performed a correlation analysis on lesion area measured by OCT at 7 days after stroke and the resulting graft survival (**Figure 6E**). The strong negative correlation between graft survival and acute lesion area disappeared in the subacute phase. However, repeating the above predictive analysis for stroke lesion size at 1 and 7 days after stroke showed a similar pattern to the previous analysis with 6/7 lesions being smaller than expected, suggesting that effects from NPC grafts are primarily conferred in the first few days after grafting (**Figure 6F-G**). Since this conferred reduction in lesion size is uncorrelated with total HA, it is possible that the early conferred effects of the NPC do not necessarily predict that there will be long-term survival of the graft in the lesion environment. This is supported by the similar lack of correlation between graft survival and separation of the brain surface at Day 7 post stroke (**Figure 6H**), as well as with OCT score (**Figure 6I**) and IHC score (**Figure 6J**). Despite a significant increase in immediate brain surface separation in NPC graft recipients (**Figure 4F.i.**), and a significant reduction in IHC score, these parameters are not correlated with long-term graft survival. As a result, despite the powerful prognostic potential of acute OCT measurements, its inferential capabilities for subacute measurements of graft efficacy cannot be effectively validated.

These data show that OCT imaging is best suited for evaluations of cell graft feasibility in acute stroke lesions, where stroke size and shape can be well characterized by OCT scattering, and the lesion environment is not yet remodeled by natural subacute wound healing processes. The ability to predict cell graft fate as a function of lesion size furthermore supports the assertion that cell graft efficacy is heavily predicated on cues from the stroke lesion environment.

## Discussion

Stroke is a leading cause of death with an urgent need for new treatments beyond the limited acute window. NPC grafting holds promise as a therapy for stimulating neural repair at the sub-acute stage, while the current gold standard for ischemic stroke treatment (tissue plasminogen activator, tPa) is limited by the need for acute administration. Here, we explored using OCT imaging to longitudinally track grafting outcomes in a photothrombotic model of stroke. OCT allows for non-invasive, intravital imaging of the mouse cortex and quickly generates volumetric images of backscattered light intensity. While this spatial data can provide useful structural information, it can also be processed further to generate angiograms and scattering maps, from which detailed quantitative information can be obtained about the cortical vasculature and the stroke lesion, respectively. Using this imaging methodology, we collected OCT data from photothrombotic strokes in the mouse cortex prior to injury and up to 2 weeks post injury, after which we conducted histological assessments on the brain tissue. In this study, we applied multiple forms of OCT post-processing as described above, which enabled us to derive angiograms, coronal profiles and scattering maps. We collected this information from mouse cortex longitudinally and quantified capillary density, brain separation from surface, and lesion size in cohorts of photothrombosis with saline injections and photothrombosis with NPC grafts. After our terminal timepoint, we showed as previously seen in a separate ischemic stroke model (Adewumi et al., 2024) that NPC grafts alter wound healing outcomes in stroke lesions by augmenting the population of wound repair astrocytes at the lesion. We also demonstrated that OCT allows for the detection of NPC graft-induced repair at lesion sites. By evaluating various correlation analyses we showed that lesion features detected by OCT correlated with total Cd13 by IHC but not other IHC-derived metrics. The findings of this study have broad implications across several areas including for: (i) use of OCT for non-invasively and longitudinally tracking stroke lesion phenotypes; (ii) the interpretation of OCT imaging data from a cell and molecular perspective; and (iii) understanding of how different methods of inducing stroke in animal models alter graft outcomes.

Widening the therapeutic window post-stroke creates a need for the longitudinal evaluation of stroke core characteristics. Typically, animal studies using histological analyses require a single time point for evaluation which limits the ability to obtain dynamic information as well as animal specific baseline information and evolution of animal-specific injury over time. Our findings demonstrate proof-of-principle that OCT can be used to track natural and treatment directed changes in wound repair and stroke lesion cores and has broad implications for the use of OCT to (i) characterize the stage of wound repair in stroke lesions in a non-invasive manner and (ii) guide treatment interventions (e.g. identify an appropriate time for grafting cells in an individual, as well as whether that individual is a suitable candidate for grafting). By extending the time window previously used for OCT imaging of photothrombotic stroke (Choi et al., 2019; Sunil et al., 2020), we were able to acquire quantitative, dynamic information about lesion area, capillary density, brain surface separation, and lesion circularity across multiple phases of wound healing. This information provided by OCT correlates to information derived from histological assessments at a single timepoint, showing that OCT is an effective and time-efficient tool for assessing cortical stroke lesions, and could be employed for the longitudinal assessment of other cortical stroke models and treatments. Historically, infarct size has been tracked in the clinic with diffusion-weighted MRI (DWI) (Moseley et al., 1990; van Everdingen et al., 1998), with measurements remaining accurate up to 14 days after stroke, well into the subacute wound healing phase (Schulz et al., 2004). A primary contribution to DWI signal intensity is the apparent diffusion coefficient (ADC) of water in the tissue, which is reduced by the edema that occurs immediately following ischemic stroke. It is reasonable that edematous tissue would exhibit increased optical scattering when imaged with OCT, with increased separation of lipid cell membranes creating a more heterogeneous distribution of scattering elements in the cortex. Additionally, the improved ability of OCT scattering for measuring lesion area compared to OCT angiograms is analogous to the well-documented “mismatch” of stroke lesion areas measured by perfusion-weighted MRI (PWI) and DWI in the clinic (Straka et al., 2010). The ability to characterize ischemic strokes longitudinally is crucial for studying subacute interventions such as cell grafts, but the incomplete understanding of molecular and cellular contributions to both DWI and OCT signal suggests a complementary need for the histological analysis of subacute stroke lesions that we performed, in both treated and untreated cases. Since the resolution of OCT imaging far exceeds that of MRI-based methods, its capabilities for multiparametric assessment of stroke lesions holds great promise for future animal studies of stroke therapeutics, especially as individual cellular contributions to the signal are further decomposed. However, OCT-based methods remain limited for stroke imaging into the subacute phase, as emergent optical scattering and increased separation of stroke lesions from the optical window led to poor signal from the lesion, as well as discrepancies between lesion assessments via OCT and IHC. Ultrasound-based imaging methods such as ultrasound localization microscopy or functional ultrasound may provide alternative future paths for subacute stroke lesion imaging, owing to ultrasound’s increased penetration depth and decoupling from optical scattering (Chavignon et al., 2022; Kılıç et al., 2020). This would allow us to explore longitudinal imaging of stroke models inaccessible with conventional optical methods (e.g. striatal stroke), where treatment outcomes from cell grafting may be enhanced.

Our findings represent significant progress toward a better understanding of the underlying cell and tissue level features that shape the outputs of OCT imaging and analysis. IHC has been extensively used by ourselves and others to characterize the cellular heterogeneity of ischemic stroke lesions, assessing angiogenesis (Krupinski et al., 1994), macrophage infiltration (Pedragosa et al., 2018), and the formation of astrocyte borders (Adewumi et al., 2024). How stroke lesion features individually contribute to OCT signal intensity and OCT scattering is not well established in the subacute phase, as imaging studies have generally focused on the acute window where treatment is most feasible. The ability to characterize both natural and treatment induced cellular remodeling of subacute stroke lesions is crucial, particularly as a primary treatment effect of NPC grafting is increased glial repair. Our IHC analysis, consistent with prior IHC-based assessments of photothrombotic stroke (Liu et al., 2017), captured the multicellularity of the subacute stroke lesion, and identified multiple potential instances of remodeling-induced scattering, particularly reactive astrocyte borders consisting of dense, interdigitating, fibrous astrocytes. This underscored a notable limitation of our OCT scattering analysis-its use as a binary classifier of ischemic and healthy tissue appeared to be not wholly compatible with subacute, remodeled stroke lesions. Among quantified cell types, only peripherally derived macrophages (Cd13) were found to be correlated with OCT measurements of lesion area. This implied that OCT measurements were most accurate for assessing large, severe lesions, but performed less consistently in smaller lesions with increased glial repair. Another notable confound is that the anisotropic contraction effect imparted by NPC grafts primarily occurred perpendicular to the imaging window (**SI** **Figure 4**). These examples of anisotropy and heterogeneity likely interfered with the application of our OCT scattering model, which was limited by the assumption that tissue scattering was consistent as a function of depth. An improved, albeit computationally expensive, scattering model would discard this assumption and estimate a scattering coefficient for every voxel in a C-scan. A three-dimensional map of tissue scattering coefficients could better characterize the layers of remodeled tissue, such as the inflamed meninges, glia limitans, Cd13-rich stroke core, and the dense reactive astrocyte borders encircling the lesion core. Such “depth-resolved” scattering models have been recently developed and tested in tissue phantoms (Almasian et al., 2015; Kübler et al., 2021), suggesting a promising transition to *in vivo* studies and eventually stroke models, following the necessary validation. Our OCT angiography methods could also be improved-as previously mentioned, we failed with OCT to detect angiogenesis within the stroke core that was detected by PECAM-1 staining, potentially due to multiple limitations-that increased surface scattering at subacute time points resulting in decreased SNR in the stroke core, or that the stroke core was populated with slow-flowing or nonfunctional vessels in which flowing RBCs did not sufficiently decorrelate to be detected by OCT angiography. Tightly constraining the initial size and severity of photothrombotic stroke lesions has been explored, and is achievable with refinements to the monitoring and control of the laser diode used (Alaverdashvili et al., 2015). Generally, the ability to reliably initiate smaller photothrombotic strokes would cap the amount of subacute inflammation interfering with our OCT imaging. A limit of the OCT angiography used in this study is the lack of quantitative information obtained about blood flow velocity and direction. To combat this, Doppler OCT has been used extensively in acute models of ischemic stroke (Choi et al., 2019; Srinivasan et al., 2013; Yu et al., 2010). Doppler OCT of the infarct and peri-infarct region could help improve our conjectures about blood flow within subacute infarcts, in order to better distinguish whether blood vessels within the stroke core are functional.

Stroke models in animal studies are typically induced by middle cerebral artery occlusion (MCAO), targeted photothrombosis (as in this study), and injection of various chemical agents or tissue degrading enzymes (Adewumi et al., 2024; Barthels and Das, 2020; O’Shea et al., 2020; Sunil et al., 2020; Van Slooten et al., 2015). While all these different methodologies of stroke induction result in damage to neural tissue there are notable differences in key lesion features such as the presence of thrombus, the density of astrocyte borders, and the composition of the peripherally immune and stromal cells that invade from the periphery (Adewumi et al., 2024; O’Shea et al., 2020). In our previous studies, we have grafted NPC and other neural cells into ischemic and hemorrhagic strokes created by injections of L-NIO (N5-(1-iminoethyl))-L-ornithine) and Collagenase 1 solutions respectively. We identified that acute lesion environments created by L-NIO provide some as yet to be identified trophic support to grafted cells such that, irrespective of the pre-graft transcriptional state (NPC, immature astrocytes, or quiescent astrocytes), grafts survive better at L-NIO lesions than all other environments, including healthy striatum and hemorrhagic stroke (Adewumi et al., 2024). While we showed here that grafts into photothrombotic strokes survive in modest numbers and direct lesion remodeling, the ischemic stroke lesions created by this particular model ultimately lack the trophic support for cell grafts that is present in L-NIO lesions (**SI** **Figure 5**). The survival of grafted cells scaled with initial lesion size, but it is notable that the remodeling detected at 2 weeks was not dependent on the total number of surviving cells. This suggests that the cause of remodeling may not only depend on the number of surviving cells but paracrine signaling to host glial cells by these surviving cells (Azevedo-Pereira et al., 2023; Rust et al., 2024). Future studies should focus on identifying the specific cellular and molecular elements impacting survival of grafted cells that are present in L-NIO stroke lesions but are notably absent in other types of injuries (Choi et al., 2016; Hou et al., 2017) as well as explore the use of biomaterial-based systems or localized, controlled molecular delivery to aid survival and promote transient proliferation of grafted cells.

In conclusion, our findings demonstrate proof-of-principle that OCT can be used to track natural and treatment directed changes in wound repair at stroke lesion cores. Furthermore, we have demonstrated specific use cases for OCT to (i) longitudinally and non-invasively characterize the stage of wound repair in stroke lesions and (ii) identify the viability of a cell grafting intervention based on OCT scattering-based measurements of acute stroke lesion size.

## Methods

### Instrumentation

For OCT imaging, a spectral domain OCT system was used (1310 nm center wavelength, bandwidth 170 nm, Thorlabs), with 5X and 10x objectives (Mitutoyo).

For initiation of photothrombosis, a 520 nm laser diode was used (L520P50, 50 mW, Thorlabs).

For monitoring of cerebral blood flow during photothrombosis, a laser speckle contrast imaging (LSCI) system was used, consisting of a 785 nm laser diode (LP785-SAV50, Thorlabs), and CMOS camera (Basler acA2040-90 μm NIR, 2048 × 2048 pixels, 5.5 × 5.5 μm pixel size). System design and real-time speckle contrast calculations were consistent with previous studies (Sunil et al., 2020).

### Derivation and Culture of Neural Progenitor Cells

Neural Progenitor Cells (NPC) were derived from female mouse embryonic stem cells (mESC) that express the hemagglutinin (HA) epitope tag (RiboTag) on modified ribosomal protein L22 (Rpl22) through neural induction and expansion as described previously (O’Shea et al., 2022). Briefly put, mESC were cultured on 0.1 % gelatin coated flask in mESC media supplemented with Leukemia inhibitory Factor (LIF). Differentiation was initiated by the removal of LIF to form embryoid bodies and then the introduction of differentiation media to induce neural cells. The neurally induced cells were expanded in neural expansion media supplemented with Epidermal Growth Factor and Fibroblast Growth Factor. NPC were passaged every 3-4 days and maintained at a passage number of less than 30.

### Surgery Procedures

#### Animals

All *in vivo* experiments and surgical procedures were performed by approved Boston University IACUC Protocol (PROTO201800520). Male and Female C57BL/6 mice (JAX#000664) were received and used for experiments between 4 and 6 months of age. Animal Housing at Boston University Animal Facility includes a 12 hour light/dark cycle, controlled temperature and humidity. Dexamethasone (4.8mg/kg), as anti-inflammatory was administered intraperitoneally 2 hours prior to the surgery, and Buprenorphine SR (slow release) 1mg/kg and Meloxicam 5mg/kg as analgesics were administered subcutaneously 1 hours prior to the surgery. For surgery, either ketamine/xylazine 100mg/10mg/kg was administered peritoneally as anesthesia, or 3% Isoflurane in oxygen for induction and 1% Isoflurane in oxygen to maintain throughout the surgery. During surgery, the mouse body temperature was kept at 37°C with the heating pad.

#### Craniotomy and window placement

Toe pinch and respiratory rate observation was monitored throughout the surgery to ascertain the depth of the anesthesia. Hair removal cream was used to remove the hair on the scalp of the mice, an incision was made and a craniotomy was performed on the left hemisphere as a 4 mm circle. Two circular glasses (3mm and 4mm) were placed on the surface of the brain and sealed with dental acrylic. An aluminum head post was also fixed using dental acrylic to allow for placement during the imaging process. The mice were allowed to recover for 2 weeks.

#### Photothrombotic Stroke

Photothrombosis was performed on a distal pial branch of the middle cerebral artery (MCA), as described previously (Sunil et al., 2020). Briefly, mice were head-fixed, and baseline LSCI was acquired for 10 minutes. A retro orbital injection of Rose Bengal (100 μl, 15 mg/ml in saline) was performed, and a kinematic mirror was used to steer the laser beam to the target vessel for 10 minutes, while monitoring the size visually to match other animals. If the infarcted area was too small, we targeted collaterals as needed. The photothrombotic laser was turned off once the target vessel was successfully occluded, as evidenced by its disappearance in LSCI images. The occlusion was monitored for one hour; if the vessel reperfused and blood flow was restored, the laser was turned on again, until subsequent vessel occlusion.

#### Cell injection

NPC were trypsinized and concentrated by centrifugation. EGF and FGF were added to the cell pellet before injection. Cells were back loaded into borosilicate pulled glass micropipettes. Four 1 uL (200k cell/uL) of cells were injected into photothrombosed mouse cortex at 0.15 uL/minute. Injections were made to target quadrant edges of the infected area, -1mm D/V. During the injections, the micropipette was moved to -0.7 mm at half volume and allowed to rest at -0.5 mm D/V for 4 minutes and slowly removed thereafter.

#### Transcardial Perfusions

At experimental endpoints, mice were overdosed with isoflurane for deep anesthesia. Cold Phosphate Buffered saline (PBS) was transcardially perfused for 2 minutes followed by 6 minutes of perfusion with freshly made 4% paraformaldehyde (from 32%) at a rate of 7mL/min. Brains were dissected and post-fixed in 4% paraformaldehyde for 6-8 hours and then transferred into 30% sucrose solution and a secondary 30% sucrose solution 24 hours later for at least 72 hours.

#### Histology and immunochemistry (IHC)

Mouse brain tissue was sliced coronally into 40μm sections using MICROM HM 525 or Leica CM1950 Cryostat. Sliced sections were stored free floating in 1X Tris Buffered Saline (TBS) supplemented with 0.001% sodium azide at 4°C. Tissue sections were stained free floating through antigen retrieval (by 1 N HCl) and several washes in 1X TBS. Brain sections were blocked and permeabilized for an hour in 1X TBS with 5% Normal donkey serum. Primary (overnight) and secondary (2 hours) antibodies were used, and nuclei were stained using DAPI (10 min). Prolong Gold mounting medium was used to coverslip the mounted brain sections on slides. Slides were allowed to dry overnight before imaging. The stained sections were imaged using IX83 Olympus Microscope.

#### Quantitative Analysis of IHC

All quantitative analyses of IHC were performed using ImageJ/FIJI. Radial angle profile intensity traces were conducted using the radial angle profile function, to derive the traces and added to collect area under the curve. IHC axial lesion was calculated by averaging the length of the neuron devoid area across 3 brain sections using plot profile function. Lesion anisotropy was derived from the ratio of the NeuN-negative profile in the X and Z direction of the largest brain section per animal.

#### Statistical analysis

GraphPad Prism 10 was utilized to construct graphs, and statistical evaluations were performed by running Student’s t-tests unless otherwise specified. Principal component analysis was performed using XLStat.

#### Optical coherence tomography image acquisition & initial processing

##### OCT angiograms

Data was acquired for the entire optical window (5x/0.14 NA objective, 3mm x 3mm FOV) and 2 regional ROIs (10x/0.28 NA objective, 1.5 mm × 1.5 mm FOV) per mouse, with scans of 1000 × 1000 pixels yielding pixel sizes of 3 µm^2^ and 1.5 µm^2^, respectively. B-scans were repeated twice and subtracted to obtain OCTA B-scans, then rastered into an OCTA C-scan. OCTA C-scans were repeated 2 times (low NA) or 5 times (high NA) and averaged to create angiograms.

##### OCT scattering maps

Data was obtained with 5x objective over 3 mm × 3 mm field of view. Scans of 1000 × 1000 pixels yielded a pixel size of 3 µm^2^. C-scans were repeated 2 times and averaged prior to postprocessing. Speckle noise was minimized by each x-y-plane by downsampling by a factor of 2 and applying bicubic interpolation. OCT attenuation, or the depth-dependent decay of OCT signal, was obtained by fitting a first order polynomial to the natural logarithm of the data in each a-line. Fits were originally performed at 200 to 500 µm beneath the tissue surface, which was obtained by manually fitting a polysurface in MATLAB. Further adjustments to the fit depth were required to account for inconsistent meningeal swelling between mice.

#### OCT postprocessing and quantitative analysis

##### OCT angiograms

For each mouse and time point, within a 10x angiogram MIPz, rectangular ROIs were drawn and classified as either stroked or healthy after superimposition of the lesion boundary from the corresponding OCT scattering map. ROIs were then analyzed for total capillary network length using the open-source application Angiotool (Zudaire et al., 2011). The reported total capillary network length for each ROI was divided by the ROI area to determine the network density.

##### Coronal Profiles

To generate coronal profiles for analysis, the raw signal from individual B-scans was log-transformed, then binned with intensity values from -3 to 2 to improve legibility. For measuring separation of the meninges and brain surface from the optical window, B-scans were serially sampled across the field of view, 20μm apart, and the maximum distance between visible interfaces was calculated using FIJI. The meninges boundary was defined as the first visible interface beneath the optical window, whereas the brain surface boundary was defined as the deepest visible interface beneath the optical window.

##### OCT scattering maps

Stroke lesion area in the axial plane was extracted from OCT scattering maps using a custom MATLAB script. Briefly, an active contour algorithm was used, in which an initial curve was drawn around the extent of the visible lesion, and iteratively evolved to the lesion boundaries, i.e. the border between large negative pixel values and small negative pixel values. The lesion boundary was used to generate a binary mask of the lesion, which was used to calculate the lesion area.

## Acknowledgements

This work was supported by internal Boston University start-up funds (T.M.O), the Craig H. Neilsen Foundation (732495 to T.M.O.), a BU NPC Core Seed Award (T.M.O.), and the NIH National Institute of Neurological Disorders and Stroke (R21NS128821 to T.M.O.). H.O.A was funded by a Boston University Biomedical Engineering Distinguished Fellowship and Faculty for the Future from Schlumberger Foundation. M.G.S was funded by the Boston University Biomedical Engineering Distinguished Fellowship and Deans Catalyst Award, College of Engineering, Boston University.

This research was supported by the Biomedical Engineering Core Facilities at Boston University. A special thanks to Xin Brown and the Biointerface Technologies (BIT) core, as well as the Micro and Nano Imaging (MNI) core at BU. We thank the Boston University Neurophotonics Center for the surgical suite, photothrombosis laser, LSCI, and OCT systems used in this work.

## Data availability

All data generated for this study are included in the main and supplementary figures. For all quantitative figures, the results of statistical tests are provided with the paper. Files of source data of individual values and other data that support the findings of this study are available on reasonable request from the corresponding author.

## Credit authorship contribution statement

**Honour O Adewumi:** Data curation, Methodology, Formal analysis, Investigation, Project administration, Supervision, Visualization, Writing – original draft, Writing – review and editing **Matthew G Simkulet:** Data curation, Methodology, Formal analysis, Investigation, Software, Visualization, Writing – original draft, Writing – review and editing **Gülce Küreli:** Data curation, Methodology, Formal analysis, Investigation, Project administration, Supervision, Writing – review and editing

**John T Giblin:** Methodology, Supervision, Software, Writing – review and editing

**Arnaldo Bisbal Lopez:** Investigation

**Şefik Evren Erdener:** Conceptualization, Writing – review and editing

**John Jiang:** Investigation

**David A Boas:** Conceptualization, Methodology, Funding acquisition, Project administration, Supervision, Resources, Writing – review and editing

**Timothy M O’Shea:** Conceptualization, Methodology, Funding acquisition, Project administration, Supervision, Resources, Visualization, Writing – original draft, Writing – review and editing

## References

1. Adewumi, H.O., Berniac, G.I., McCarthy, E.A., O’Shea, T.M., 2024. Ischemic and hemorrhagic stroke lesion environments differentially alter the glia repair potential of neural progenitor cell and immature astrocyte grafts. Exp. Neurol. 374, 114692. 10.1016/j.expneurol.2024.114692

2. Alaverdashvili, M., Paterson, P.G., Bradley, M.P., 2015. Laser system refinements to reduce variability in infarct size in the rat photothrombotic stroke model. J. Neurosci. Methods 247, 58–66. 10.1016/j.jneumeth.2015.03.029

3. Almasian, M., Bosschaart, N., Leeuwen, T.G. van, Faber, D.J., 2015. Validation of quantitative attenuation and backscattering coefficient measurements by optical coherence tomography in the concentration-dependent and multiple scattering regime. J. Biomed. Opt. 20, 121314. 10.1117/1.JBO.20.12.121314

4. Azevedo-Pereira, R.L., Manley, N.C., Dong, C., Zhang, Y., Lee, A.G., Zatulovskaia, Y., Gupta, V., Vu, J., Han, S., Berry, J.E., Bliss, T.M., Steinberg, G.K., 2023. Decoding the molecular crosstalk between grafted stem cells and the stroke-injured brain. Cell Rep. 42. 10.1016/j.celrep.2023.112353

5. Barthels, D., Das, H., 2020. Current advances in ischemic stroke research and therapies. Biochim. Biophys. Acta BBA - Mol. Basis Dis., Stem Cells and Their Applications to Human Diseases 1866, 165260. 10.1016/j.bbadis.2018.09.012

6. Berry, M., Maxwell, W.L., Logan, A., Mathewson, A., McConnell, P., Ashhurst, D.E., Thomas, G.H., 1983. Deposition of scar tissue in the central nervous system. Acta Neurochir Suppl Wien. 10.1007/978-3-7091-4147-2_3

7. Burda, J.E., Sofroniew, M.V., 2014. Reactive gliosis and the multicellular response to CNS damage and disease. Neuron. 10.1016/j.neuron.2013.12.034

8. Chavignon, A., Hingot, V., Orset, C., Vivien, D., Couture, O., 2022. 3D transcranial ultrasound localization microscopy for discrimination between ischemic and hemorrhagic stroke in early phase. Sci. Rep. 12, 14607. 10.1038/s41598-022-18025-x

9. Choi, W.J., Li, Y., Wang, R.K., 2019. Monitoring Acute Stroke Progression: Multi-Parametric OCT Imaging of Cortical Perfusion, Flow, and Tissue Scattering in a Mouse Model of Permanent Focal Ischemia. IEEE Trans. Med. Imaging 38, 1427–1437. 10.1109/TMI.2019.2895779

10. Choi, Y.-K., Urnukhsaikhan, E., Yoon, H.-H., Seo, Y.-K., Park, J.-K., 2016. Effect of human mesenchymal stem cell transplantation on cerebral ischemic volume-controlled photothrombotic mouse model. Biotechnol. J. 11, 1397–1404. 10.1002/biot.201600057

11. Dias, D.O., Kalkitsas, J., Kelahmetoglu, Y., Estrada, C.P., Tatarishvili, J., Ernst, A., Huttner, H.B., Kokaia, Z., Lindvall, O., Brundin, L., Frisén, J., Göritz, C., 2021. Pericyte-derived fibrotic scarring is conserved across diverse central nervous system lesions. Nat. Commun. 10.1038/s41467-021-25585-5

12. Feigin, V.L., Norrving, B., Mensah, G.A., 2017. Global Burden of Stroke. Circ. Res. 120, 439–448. 10.1161/CIRCRESAHA.116.308413

13. Goncalves, A., Su, E.J., Muthusamy, A., Zeitelhofer, M., Torrente, D., Nilsson, I., Protzmann, J., Fredriksson, L., Eriksson, U., Antonetti, D.A., Lawrence, D.A., 2022. Thrombolytic tPA-Induced Hemorrhagic Transformation of Ischemic Stroke is Mediated by PKCβ phosphorylation of Occludin. Blood blood.2021014958. 10.1182/blood.2021014958

14. Grysiewicz, R.A., Thomas, K., Pandey, D.K., 2008. Epidemiology of ischemic and hemorrhagic stroke: incidence, prevalence, mortality, and risk factors. Neurol. Clin. 26, 871–895, vii. 10.1016/j.ncl.2008.07.003

15. Hacke, W., Kaste, M., Bluhmki, E., Brozman, M., Dávalos, A., Guidetti, D., Larrue, V., Lees, K.R., Medeghri, Z., Machnig, T., Schneider, D., von Kummer, R., Wahlgren, N., Toni, D., 2008. Thrombolysis with Alteplase 3 to 4.5 Hours after Acute Ischemic Stroke. N. Engl. J. Med. 359, 1317–1329. 10.1056/NEJMoa0804656

16. Hou, B., Ma, J., Guo, X., Ju, F., Gao, J., Wang, D., Liu, J., Li, X., Zhang, S., Ren, H., 2017. Exogenous Neural Stem Cells Transplantation as a Potential Therapy for Photothrombotic Ischemia Stroke in Kunming Mice Model. Mol. Neurobiol. 54, 1254–1262. 10.1007/s12035-016-9740-6

17. Jickling, G.C., Liu, D., Stamova, B., Ander, B.P., Zhan, X., Lu, A., Sharp, F.R., 2014. Hemorrhagic Transformation after Ischemic Stroke in Animals and Humans. J. Cereb. Blood Flow Metab. 34, 185–199. 10.1038/jcbfm.2013.203

18. Johansson, B.B., 2000. Brain plasticity and stroke rehabilitation. The Willis lecture. Stroke 31, 223–230. 10.1161/01.str.31.1.223

19. Kılıç, K., Tang, J., Erdener, Ş.E., Sunil, S., Giblin, J.T., Lee, B.S., Postnov, D.D., Chen, A., Boas, D.A., 2020. Chronic Imaging of Mouse Brain: From Optical Systems to Functional Ultrasound. Curr. Protoc. Neurosci. 93, e98. 10.1002/cpns.98

20. Krupinski, J., Kaluza, J., Kumar, P., Kumar, S., Wang, J.M., 1994. Role of angiogenesis in patients with cerebral ischemic stroke. Stroke 25, 1794–1798. 10.1161/01.STR.25.9.1794

21. Kübler, J., Zoutenbier, V.S., Amelink, A., Fischer, J., Boer, J.F. de, 2021. Investigation of methods to extract confocal function parameters for the depth resolved determination of attenuation coefficients using OCT in intralipid samples, titanium oxide phantoms, and in vivo human retinas. Biomed. Opt. Express 12, 6814–6830. 10.1364/BOE.440574

22. Li, P., Murphy, T.H., 2008. Two-Photon Imaging during Prolonged Middle Cerebral Artery Occlusion in Mice Reveals Recovery of Dendritic Structure after Reperfusion. J. Neurosci. 28, 11970–11979. 10.1523/JNEUROSCI.3724-08.2008

23. Li, Y., He, X., Kawaguchi, R., Zhang, Y., Wang, Q., Monavarfeshani, A., Yang, Z., Chen, B., Shi, Z., Meng, H., Zhou, S., Zhu, J., Jacobi, A., Swarup, V., Popovich, P.G., Geschwind, D.H., He, Z., 2020. Microglia-organized scar-free spinal cord repair in neonatal mice. Nature. 10.1038/s41586-020-2795-6

24. Liang, H., Zhao, H., Gleichman, A., Machnicki, M., Telang, S., Tang, S., Rshtouni, M., Ruddell, J., Carmichael, S.T., 2019. Region-specific and activity-dependent regulation of SVZ neurogenesis and recovery after stroke. Proc. Natl. Acad. Sci. 116, 13621–13630. 10.1073/pnas.1811825116

25. Liu, N.-W., Ke, C.-C., Zhao, Y., Chen, Y.-A., Chan, K.-C., Tan, D.T.-W., Lee, J.-S., Chen, Y.-Y., Hsu, T.-W., Hsieh, Y.-J., Chang, C.-W., Yang, B.-H., Huang, W.-S., Liu, R.-S., 2017. Evolutional Characterization of Photochemically Induced Stroke in Rats: a Multimodality Imaging and Molecular Biological Study. Transl. Stroke Res. 8, 244–256. 10.1007/s12975-016-0512-4

26. Magid-Bernstein, J., Girard, R., Polster, S., Srinath, A., Romanos, S., Awad, I.A., Sansing, L.H., 2022. Cerebral Hemorrhage: Pathophysiology, Treatment, and Future Directions. Circ. Res. 130, 1204–1229. 10.1161/CIRCRESAHA.121.319949

27. Maxwell, W.L., Follows, R., Ashhurst, D.E., Berry, M., Boycott, B.B., 1990. The response of the cerebral hemisphere of the rat to injury. II. The neonatal rat. Philos. Trans. R. Soc. Lond. B Biol. Sci. 10.1098/rstb.1990.0122

28. Moseley, M.E., Kucharczyk, J., Mintorovitch, J., Cohen, Y., Kurhanewicz, J., Derugin, N., Asgari, H., Norman, D., 1990. Diffusion-weighted MR imaging of acute stroke: correlation with T2-weighted and magnetic susceptibility-enhanced MR imaging in cats. Am. J. Neuroradiol. 11, 423–429.

29. O’Shea, T.M., Ao, Y., Wang, S., Wollenberg, A.L., Kim, J.H., Ramos Espinoza, R.A., Czechanski, A., Reinholdt, L.G., Deming, T.J., Sofroniew, M.V., 2022. Lesion environments direct transplanted neural progenitors towards a wound repair astroglial phenotype in mice. Nat. Commun. 13, 5702. 10.1038/s41467-022-33382-x

30. O’Shea, T.M., Wollenberg, A.L., Kim, J.H., Ao, Y., Deming, T.J., Sofroniew, M.V., 2020. Foreign body responses in central nervous system mimic natural wound responses and alter biomaterial functions. Nat. Commun. 11. 10.1038/s41467-020-19906-3

31. Pedragosa, J., Salas-Perdomo, A., Gallizioli, M., Cugota, R., Miró-Mur, F., Briansó, F., Justicia, C., Pérez-Asensio, F., Marquez-Kisinousky, L., Urra, X., Gieryng, A., Kaminska, B., Chamorro, A., Planas, A.M., 2018. CNS-border associated macrophages respond to acute ischemic stroke attracting granulocytes and promoting vascular leakage. Acta Neuropathol. Commun. 6, 76. 10.1186/s40478-018-0581-6

32. Phipps, M.S., Cronin, C.A., 2020. Management of acute ischemic stroke. BMJ 368, l6983. 10.1136/bmj.l6983

33. Prust, M.L., Forman, R., Ovbiagele, B., 2024. Addressing disparities in the global epidemiology of stroke. Nat. Rev. Neurol. 20, 207–221. 10.1038/s41582-023-00921-z

34. Rosenzweig, S., Carmichael, S.T., 2013. Age-Dependent Exacerbation of White Matter Stroke Outcomes. Stroke 44, 2579–2586. 10.1161/STROKEAHA.113.001796

35. Ruikang K. Wang, Qinqin Zhang, Yuandong Li, Shaozhen Song, 2017. Optical coherence tomography angiography-based capillary velocimetry. J. Biomed. Opt. 22, 066008. 10.1117/1.JBO.22.6.066008

36. Rust, R., Nih, L.R., Liberale, L., Yin, H., El Amki, M., Ong, L.K., Zlokovic, B.V., 2024. Brain repair mechanisms after cell therapy for stroke. Brain awae204. 10.1093/brain/awae204

37. Schmitt, J.M., Knuettel, A.R., Gandjbakhche, A.H., Bonner, R.F., 1993. Optical characterization of dense tissues using low-coherence interferometry, in: Holography, Interferometry, and Optical Pattern Recognition in Biomedicine III. Presented at the Holography, Interferometry, and Optical Pattern Recognition in Biomedicine III, SPIE, pp. 197–211. 10.1117/12.155715

38. Schulz, U.G., Briley, D., Meagher, T., Molyneux, A., Rothwell, P.M., 2004. Diffusion-Weighted MRI in 300 Patients Presenting Late With Subacute Transient Ischemic Attack or Minor Stroke. Stroke 35, 2459–2465. 10.1161/01.STR.0000143455.55877.b9

39. Smith, E.E., Saposnik, G., Biessels, G.J., Doubal, F.N., Fornage, M., Gorelick, P.B., Greenberg, S.M., Higashida, R.T., Kasner, S.E., Seshadri, S., 2017. Prevention of Stroke in Patients With Silent Cerebrovascular Disease: A Scientific Statement for Healthcare Professionals From the American Heart Association/American Stroke Association. Stroke 48, e44–e71. 10.1161/STR.0000000000000116

40. Sozmen, E.G., Hinman, J.D., Carmichael, S.T., 2012. Models That Matter: White Matter Stroke Models. Neurotherapeutics 9, 349–358. 10.1007/s13311-012-0106-0

41. Srinivasan, V.J., Mandeville, E.T., Can, A., Blasi, F., Climov, M., Daneshmand, A., Lee, J.H., Yu, E., Radhakrishnan, H., Lo, E.H., Sakadžić, S., Eikermann-Haerter, K., Ayata, C., 2013. Multiparametric, Longitudinal Optical Coherence Tomography Imaging Reveals Acute Injury and Chronic Recovery in Experimental Ischemic Stroke. PLOS ONE 8, e71478. 10.1371/journal.pone.0071478

42. Steven R. Messé, Pooja Khatri, Mathew J. Reeves, Eric E. Smith, Jeffrey L. Saver, Deepak L. Bhatt, Maria V. Grau-Sepulveda, Margueritte Cox, Eric D. Peterson, Gregg C. Fonarow, Lee H. Schwamm, 2016. Why are acute ischemic stroke patients not receiving IV tPA? Neurology 87, 1565. 10.1212/WNL.0000000000003198

43. Straka, M., Albers, G.W., Bammer, R., 2010. Real-time diffusion-perfusion mismatch analysis in acute stroke. J. Magn. Reson. Imaging 32, 1024–1037. 10.1002/jmri.22338

44. Sunil, S., Erdener, S.E., Lee, B.S., Postnov, D., Tang, J., Kura, S., Cheng, X., Chen, I.A., Boas, D.A., Kilic, K., 2020. Awake chronic mouse model of targeted pial vessel occlusion via photothrombosis. Neurophotonics 7, 015005. 10.1117/1.NPh.7.1.015005

45. Sunil, S., Evren Erdener, S., Cheng, X., Kura, S., Tang, J., Jiang, J., Karrobi, K., Kılıç, K., Roblyer, D., Boas, D.A., 2021. Stroke core revealed by tissue scattering using spatial frequency domain imaging. NeuroImage Clin. 29, 102539. 10.1016/j.nicl.2020.102539

46. Tang, J., Cheng, X., Kilic, K., Devor, A., Lee, J., Boas, D.A., 2021. Imaging localized fast optical signals of neural activation with optical coherence tomography in awake mice. Opt. Lett. 46, 1744– 1747. 10.1364/OL.411897

47. van Everdingen, K.J., van der Grond, J., Kappelle, L.J., Ramos, L.M.P., Mali, W.P.T.M., 1998. Diffusion-Weighted Magnetic Resonance Imaging in Acute Stroke. Stroke 29, 1783–1790. 10.1161/01.STR.29.9.1783

48. Van Slooten, A.R., Sun, Y., Clarkson, A.N., Connor, B.J., 2015. l-NIO as a novel mechanism for inducing focal cerebral ischemia in the adult rat brain. J. Neurosci. Methods 245, 44–57. 10.1016/j.jneumeth.2015.02.022

49. Wang, R.K., Jacques, S.L., Ma, Z., Hurst, S., Hanson, S.R., Gruber, A., 2007. Three dimensional optical angiography. Opt. Express 15, 4083–4097. 10.1364/OE.15.004083

50. Yaghi, S., Willey, J.Z., Cucchiara, B., Goldstein, J.N., Gonzales, N.R., Khatri, P., Kim, L.J., Mayer, S.A., Sheth, K.N., Schwamm, L.H., 2017. Treatment and Outcome of Hemorrhagic Transformation After Intravenous Alteplase in Acute Ischemic Stroke: A Scientific Statement for Healthcare Professionals From the American Heart Association/American Stroke Association. Stroke 48. 10.1161/STR.0000000000000152

51. Yu, L., Nguyen, E., Liu, G., Choi, B., Chen, Z., 2010. Spectral Doppler optical coherence tomography imaging of localized ischemic stroke in a mouse model. J. Biomed. Opt. 15, 066006. 10.1117/1.3505016

52. Zhao, L.-R., Willing, A., 2018. Enhancing endogenous capacity to repair a stroke-damaged brain: An evolving field for stroke research. Neurobiol. Stroke Adv. Chall. Future Dir. 163–164, 5–26. 10.1016/j.pneurobio.2018.01.004

53. Zudaire, E., Gambardella, L., Kurcz, C., Vermeren, S., 2011. A Computational Tool for Quantitative Analysis of Vascular Networks. PLOS ONE 6, e27385. 10.1371/journal.pone.0027385

