## Supplementary Information for "Optical coherence tomography enables longitudinal evaluation of cell graft-directed remodeling in stroke lesions"

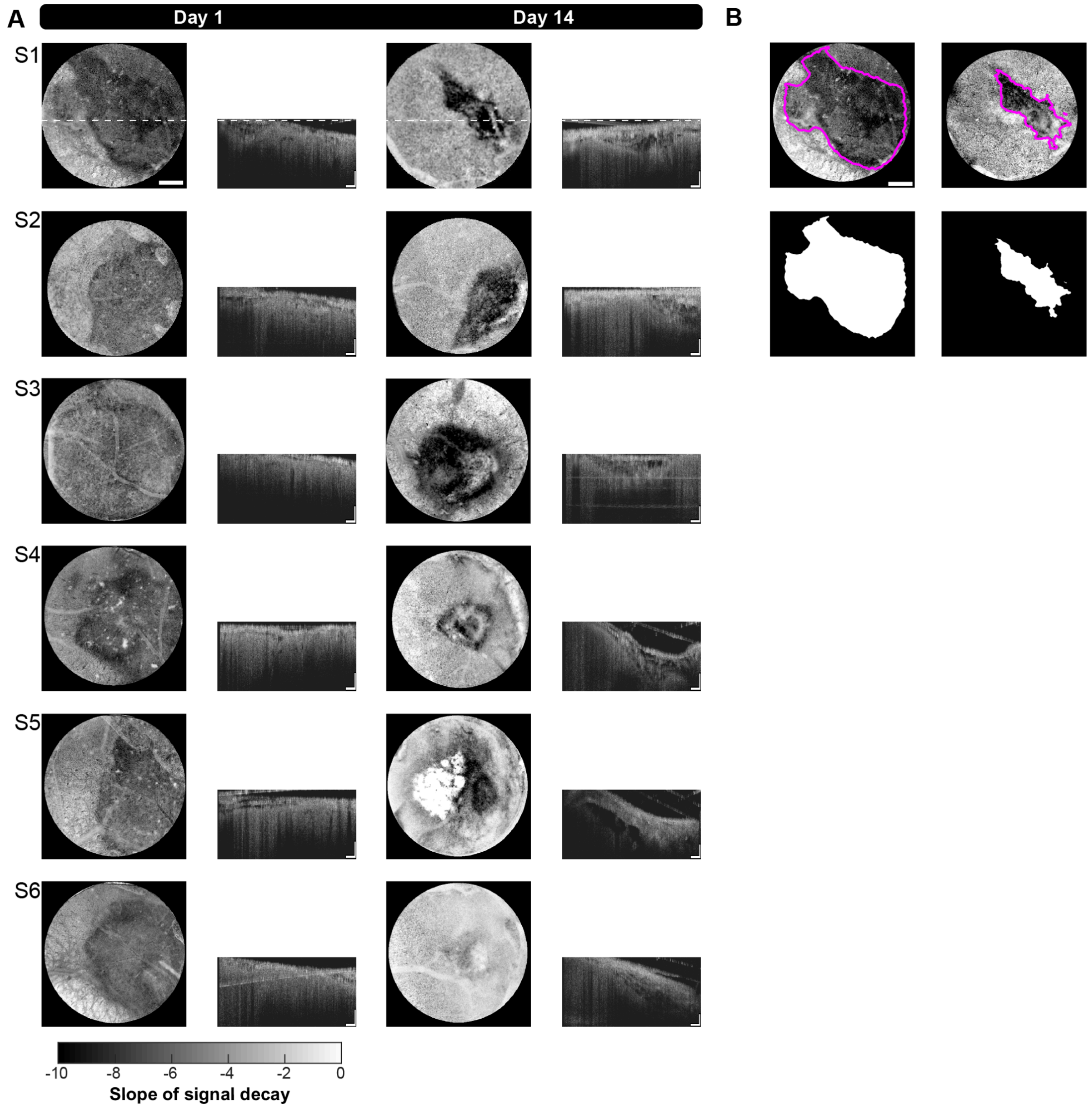


**SI Figure 1: Saline stroke core longitudinal imaging and segmentation. A.** *Left:* OCT scattering maps acquired at 1 day and 14 days after stroke for all saline control mice. Scale bar = 500μm. *Right:* Example B-scans from the coronal plane, as identified by the dotted line at the top. Scale bars = 200μm. **B.** Example segmentation of stroke cores with resulting masks. Scale bar = 500μm.

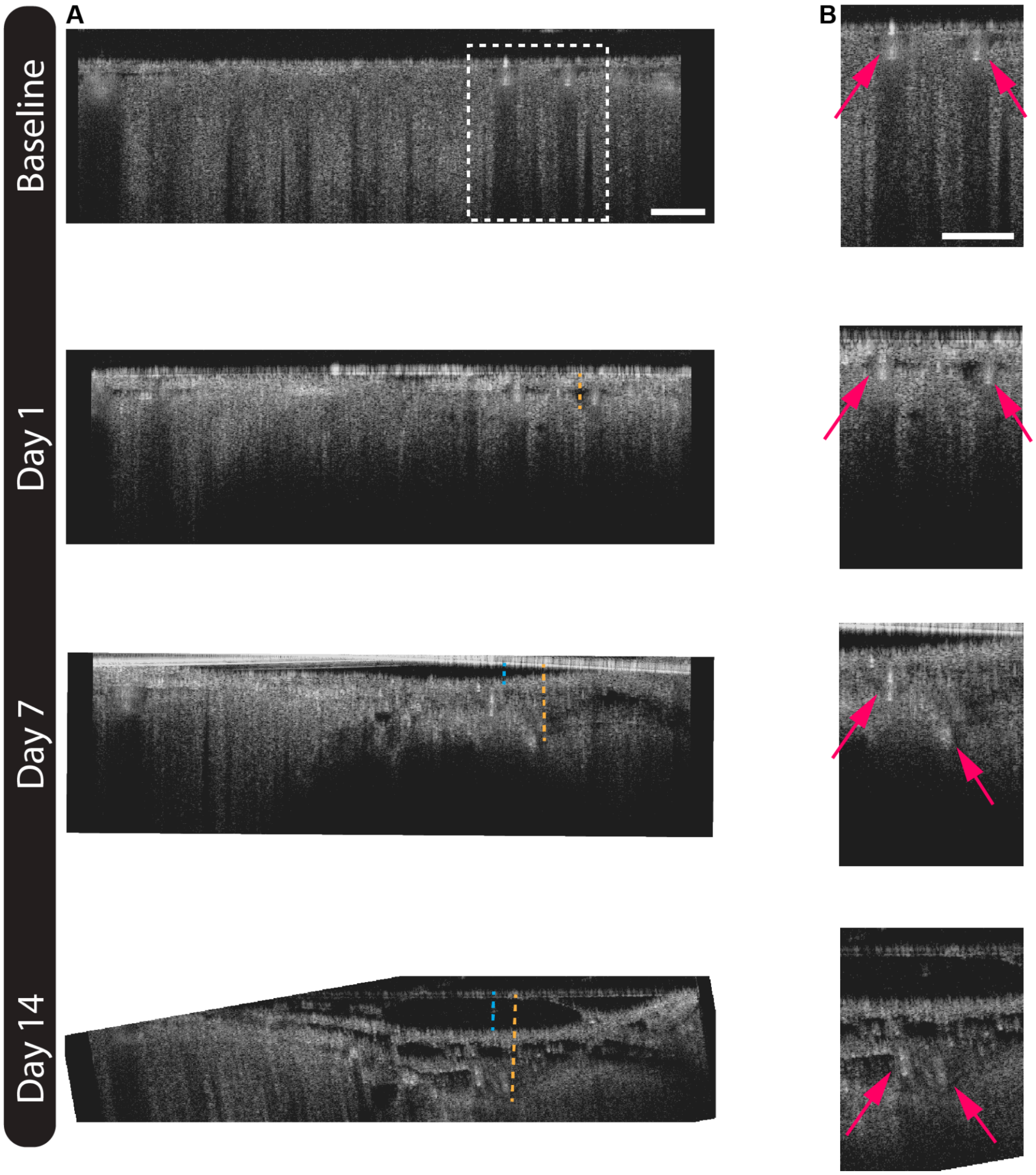


**SI Figure 2: Identification of structural features in coronal profiles. A.** Sample coronal profiles demonstrating increased cortical and meningeal separation from the imaging window. Blue dash = meninges, i.e. the first observable interface beneath the window. Orange dash = cortical surface, i.e. the deepest observable interface. Scale bar = 250µm. **B.** Inset from Part A. Magenta arrows denote major pial vessels. Scale bar = 250µm.

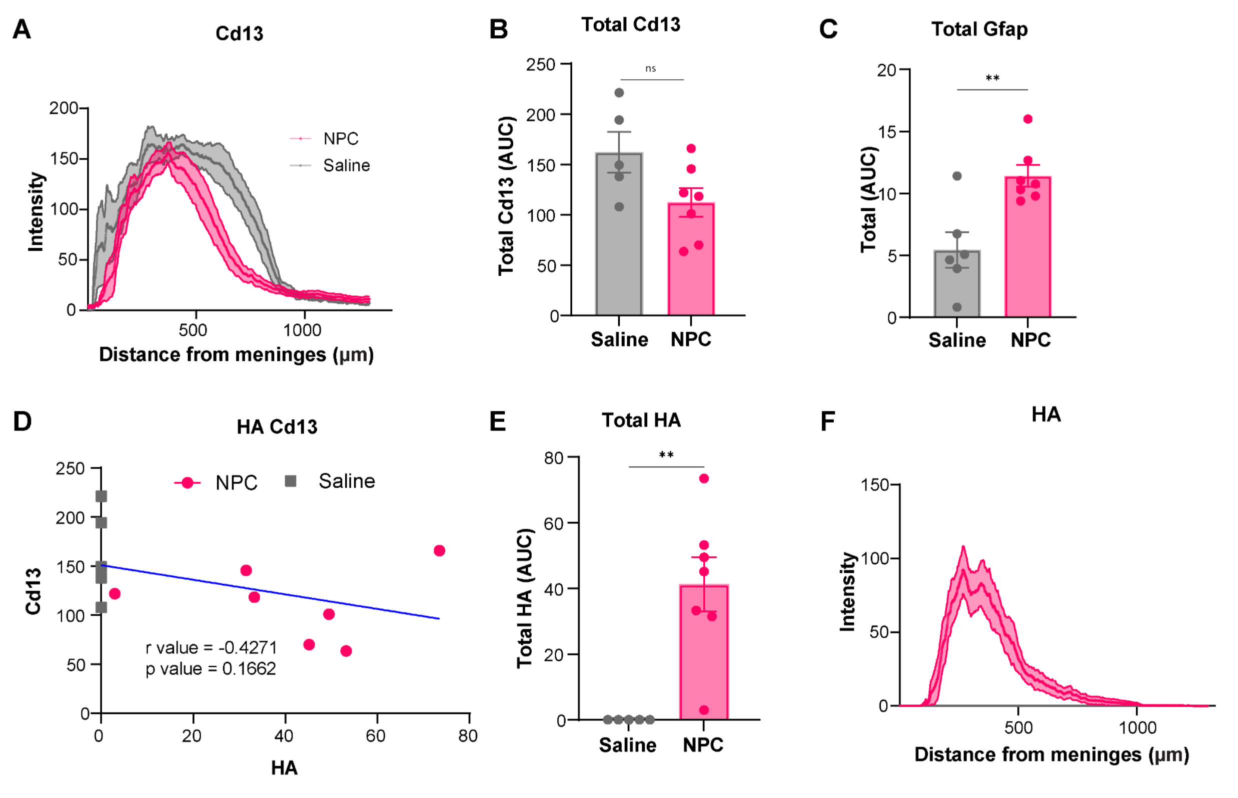


**SI Figure 3:** Quantifications from immunohistochemistry. **A**. Cd13 trace of NPC and saline groups radially from meninges into the cortex. **B**. Total area under the curve of Cd13 trace (Student's t test;ns=not significant ). **C**.Total area under the curve of Gfap trace (Student's t test; p value-0.0075).**D**. Correlation graph of total HA (Cell grafts) and total Cd13 (r value: -0.4271, p value: 0.1662). **E**. Total area under the curve of HA trace (Student's t test; p value-0.0020). **F**. Cd13 trace of NPC and saline groups radially from meninges into the cortex.

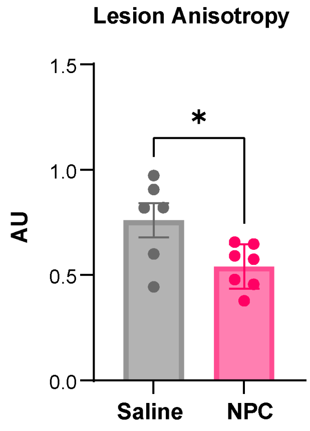


**SI Figure 4:** Stroke lesion anisotropy, as an aspect ratio, as measured by IHC. *P=0.0272, student's t-test.

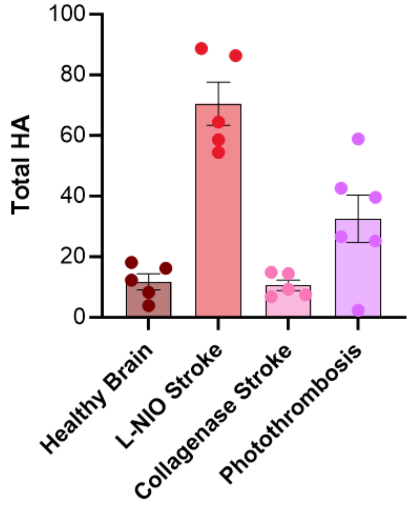


**SI Figure 5:** Immunohistochemistry of multiple stroke lesion types and total surviving NPC grafts.

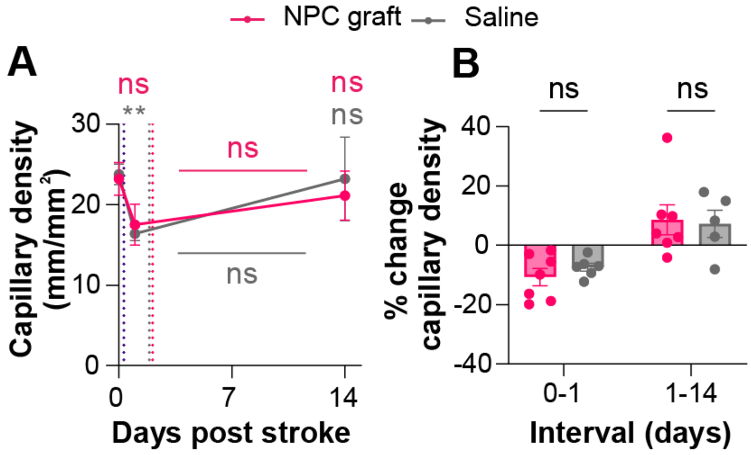


**SI Figure 6: Capillary density measured outside stroke core. A.** Longitudinal change in capillary density outside the stroke core, (mm capillary/mm$^2$ area), measured from intracortical OCTA MIPz acquired at 10x magnification and segmented as described. Not significant (ns), **P = 0.0061, mixed-effects analysis with Tukey multiple comparisons test. Colored comparisons are to the corresponding group's Day 1 or as otherwise indicated. Line plot shows mean ± s.e.m., for N = 7 NPC graft mice and N = 6 saline mice, or fewer if a given mouse did not have 10x images overlapping healthy tissue. **B.** % Change in capillary density across the given interval. Not significant (ns) and ***P < 0.0001, mixed-effects analysis with uncorrected Fisher's LSD. Bar chart shows mean ± s.e.m., with individual data points representing N = 7 NPC graft mice or N = 6 saline mice, or fewer where indicated.

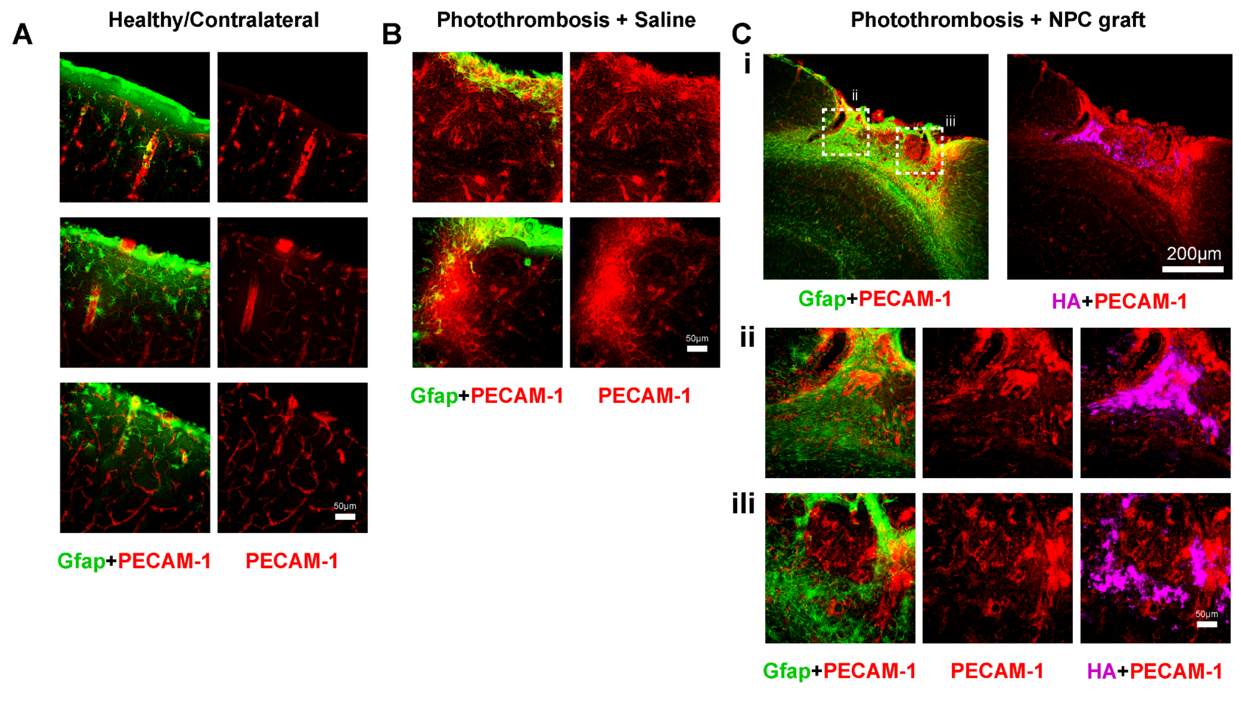


**SI Figure 7:** Immunohistochemistry of stroke lesions and healthy contralateral section. **A**. overview image of coronal brain section of healthy brain cortex. **B**. Photothromosed and saline-injected cortex. **C**. Phothothromobisis with NPC grafts at 2 days post-stroke. All showing stains for Gfap (astrocytes), PECAM-1 (vasculature), and HA (NPC grafts).

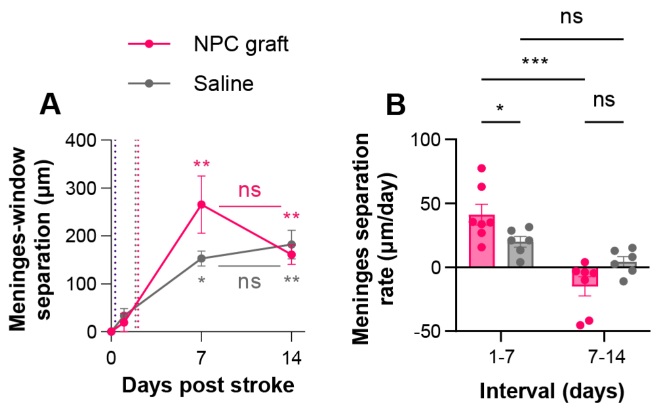


**SI Figure 8: Longitudinal separation between meninges and optical window. A.** Longitudinal change in separation between the meninges and optical window, measured from coronal profiles. Not significant (ns), *P = 0.0106, **P < 0.0063, repeated measures two-way ANOVA with Tukey multiple comparisons test. Colored comparisons are to the corresponding group's Day 1 or as otherwise indicated. Line plot shows mean ± s.e.m., for N = 7 NPC graft mice and N = 6 saline mice. **B.** Separation rate of meninges along the specified intervals. Not significant (ns), *P = 0.033, ***P = 0.0007, repeated measures two-way ANOVA with Tukey multiple comparisons test. Bar chart shows mean ± s.e.m., with individual data points representing N = 7 NPC graft mice or N = 6 saline mice, or fewer where indicated.

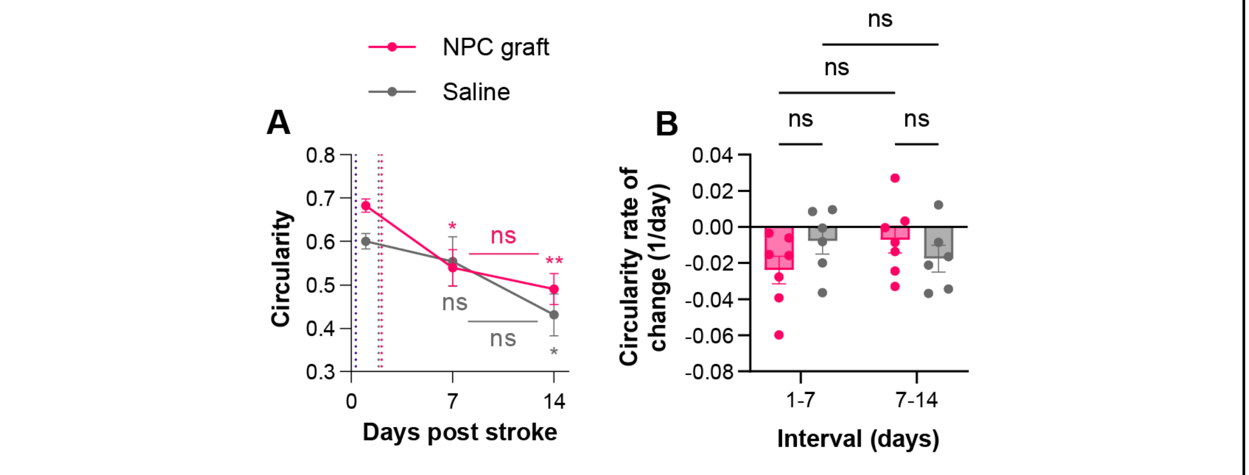


**SI Figure 9: Longitudinal change in lesion circularity derived from OCT scattering maps. A.** Longitudinal change in circularity of stroke lesions, measured from stroke core masks segmented from OCT scattering maps. Not significant (ns), $^{*}P < 0.0455$, $^{**}P = 0.0039$, repeated measures two-way ANOVA with Tukey multiple comparisons test. Colored comparisons are to the corresponding group's Day 1 or as otherwise indicated. Line plot shows mean ± s.e.m., for N = 7 NPC graft mice and N = 6 saline mice. **B.** Rate of change of circularity along the specified intervals between imaging sessions. Not significant (ns), repeated measures two-way ANOVA with Tukey multiple comparisons test. Bar chart shows mean ± s.e.m., with individual data points representing N = 7 NPC graft mice or N = 6 saline mice, or fewer where indicated.

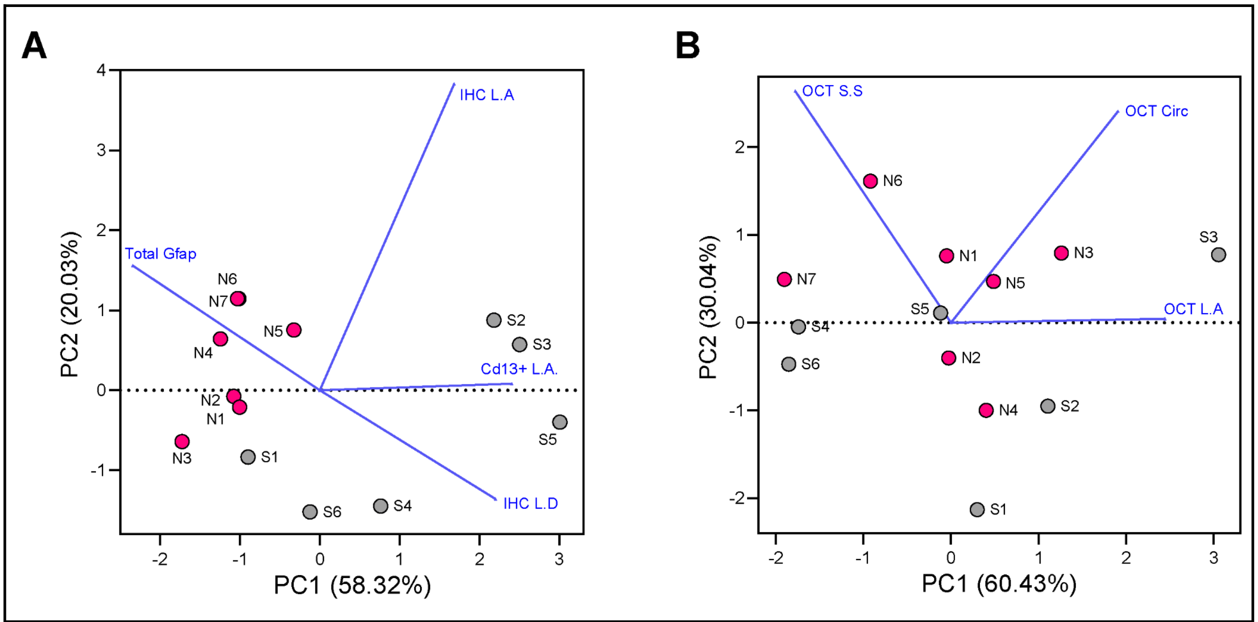


**SI Figure 10: Principal Component Analysis on OCT and IHC Parameters. A.** PCA for IHC Parameter (Total Gfap, IHC Lesion Area (L.A.), Cd13 positive Lesion area, IHC Lesion Depth). B. PCA for OCT Parameter (Surface Separation (S.S.), Circularity (Circ), Lesion area (L.A.).
